# Synergy and allostery in ligand binding by HIV-1 Nef

**DOI:** 10.1101/2020.11.05.369645

**Authors:** Abdullah Aldehaiman, Afaque A. Momin, Audrey Restouin, Luyao Wang, Xiaoli Shi, Safia Aljedani, Sandrine Opi, Adrien Lugari, Umar F. Shahul Hameed, Luc Ponchon, Xavier Morelli, Mingdong Huang, Christian Dumas, Yves Collette, Stefan T. Arold

## Abstract

The Nef protein of human and simian immunodeficiency viruses (HIV and SIV, respectively) boosts viral pathogenicity through its interactions with host cell proteins. Nef has a folded core domain and large flexible regions, each carrying several protein interaction sites. By combining the polyvalency intrinsic to unstructured regions with the binding selectivity and strength of a 3D folded domain, Nef can bind to many different host cell proteins, perturbing their cellular functions. For example, the combination of a linear proline-rich motif and a hydrophobic core domain surface allows Nef to increase affinity and selectivity for particular Src family SH3 domains. Here we investigated whether the interplay between Nef’s flexible regions and its core domain can allosterically influence ligand selection. We found that the flexible regions can bind back to the core domain in different ways, producing distinct conformational states that alter the SH3 domain selectivity and availability of Nef’s functional motifs. The resulting cross-talk might help synergising certain subsets of ligands while excluding others, promoting functionally coherent Nef-bound protein ensembles. Further, we combined proteomic and bioinformatic analyses to identify human proteins that select SH3 domains in the same way as does Nef. We found that only 2–3% of clones from a whole human fetal library displayed a Nef-like SH3 selectivity. However, in most cases this selectivity appears to be achieved by a canonical linear interaction rather than a Nef-like ‘tertiary’ interaction. This analysis suggests that Nef’s SH3 recognition surface has no (or marginally few) cellular counterparts, validating the Nef tertiary binding surface as a promising unique drug target.

## Introduction

Nef is an accessory protein important for enhancing the virulence and pathogenesis of human and simian immunodeficiency viruses (HIV and SIV respectively) (1,2). The absence of a functional Nef protein results in HIV viruses that fail to cause AIDS in infected persons (3–6). Hence, Nef is considered a promising drug target.

Nef’s role in disease progression results from its capacity to perturb several host cell functions. Nef alters the protein composition of the host cell surface, mostly by downregulating transmembrane proteins involved in immune signalling and viral entry (including CD3, CD4, CD8, CD28, CXCR4, CCR5, SERINC3, SERINC5 and antigen-loaded MHC-II) [reviewed in (7)]. However, Nef also upregulates the surface expression of other factors such as TNF, DC-SIGN and LIGHT (8,9). Other functions of Nef include the promotion of T cell activation (10,11) and lymphocyte chemotaxis (12). Nef also perturbs intracellular signalling, in particular through its interactions with cytoplasmic kinases (including members of the Src and Tec/Bt kinase families, PAK2, and the PI3 kinase) (13–18). Collectively, these actions promote immune evasion of infected cells as well as enhancement of viral release, spreading and infectivity.

Nef is a non-catalytic protein and all its effects are caused by its interactions with a multitude of cellular proteins. The molecular bases for a subset of these interactions have been determined. Initial NMR and crystallographic studies showed that Nef consists of a folded core domain (residues 71-206; herein we use the numbering of the LAI isolate unless stated otherwise), a myristoylated N-terminal flexible arm (residues 1-70) and a central core loop (residues 148-179)(**Fig 1A**) (19–23). Both the core domain and flexible regions contain important ligand binding sites (1,7).

**Fig 1.**
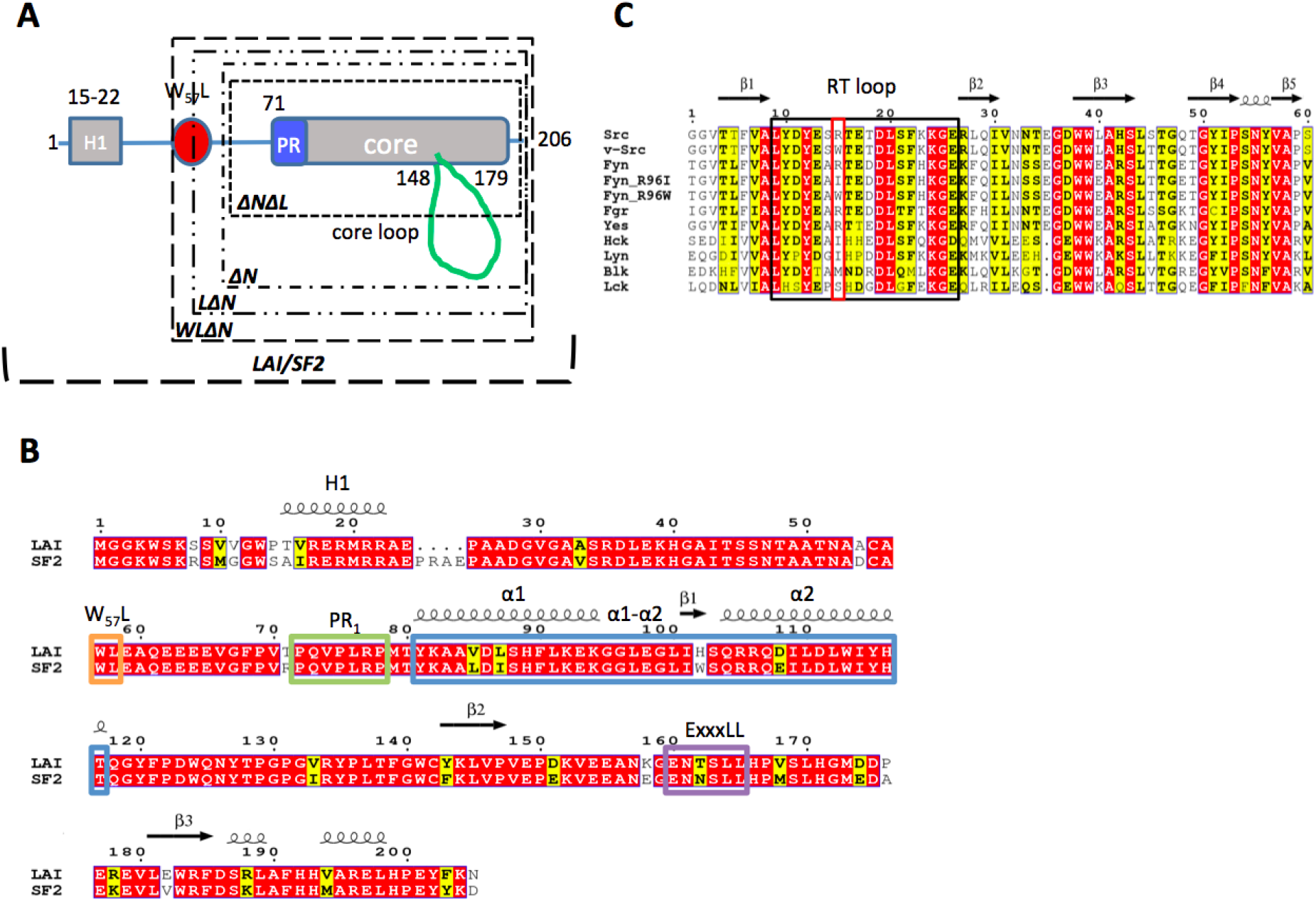
Schematic overview of the constructs used. **A)** Schematic drawing of Nef constructs used where key residues such as W_57_L and the proline rich region (PR, including the P_72_xxPxR motif), are shown. **B)** Sequence alignment between Nef LAI/SF2 where key motifs discussed in the paper are highlighted: H1 (brown), W_57_L (orange), PR (green), α1-α2 (blue), and E_160_xxxLL endocytosis motif (purple). **C)** Sequence alignment of all SH3 domains discussed. The RT loop (black) and the ‘R’ position within (red) are highlighted. In both sequence alignments, residues with similar physico-chemical properties are written with black bold letters on yellow background. Conserved residues are highlighted in red.

Subsequent structural studies revealed the molecular determinants for Nef’s capacity to perturb Src kinases by tightly interacting with their SH3 domains (20,22,24). In addition to its canonical ‘linear’ proline-rich motif P_72_xxPxR (where x is any amino acid), Nef uses an extended surface provided by its folded core to bind to Src family SH3 domains. The resulting ‘tertiary’ binding mode increases the interaction surface, and hence affinity and selectivity, compared to the canonical linear interaction between proline-rich motifs and SH3 domains (1,20,22). One particular key residue from the SH3 domain, located in the arginine (‘R’) position of the so-called arginine-threonine (RT) loop (residue number 96 in Fyn), was shown to centrally affect the binding affinity by interacting with a pocket on the Nef core surface (25). This pocket is hydrophobic in HIV-1 Nef, and more polar/charged in SIV Nef. Consequently, HIV-1 Nef associates more tightly with kinases whose SH3 domain have a hydrophobic residue in the ‘R’ position, whereas SIV Nef prefers charged or polar residues (20,26). However, additional effects were noted, such as the extent of the stabilising hydrogen bond network of the SH3 RT loop (27). The RT loop needs to change conformation to adapt to the Nef tertiary surface, and hence a more extensive hydrogen bond network (such as seen in Fyn SH3) entails a higher enthalpic cost for binding then intrinsically more flexible RT loops (case of Hck). SH3 binding by Nef activates the Src family kinases Hck by outcompeting the intramolecular interactions that maintain the inactive ‘closed’ kinase conformation (28,29). SH3-dependent activation by Nef was also observed for Lyn and c-Src in a *Saccharomyces cerevisiae* assay (30), but not in an *in vivo* transformation assay (31). The former assay also failed to show activation of Fgr, Fyn, Lck, or Yes by Nef. Hence, although a role of Nef-targeted kinases in HIV replication, infectivity, MHC-1 downregulation and pro-inflammatory vesicle release was observed (16,17,32,33), the molecular mechanism for it remains to be fully elucidated.

A series of structural studies also elucidated how Nef molecules hijack the host cell trafficking machinery by serving as an adaptor between AP1 and MHC-I molecules or between AP2 and CD4 (7,34–37). Akin to its interactions with Src family SH3 domains, Nef uses its core domain in addition to the linear recognition motifs (located on Nef’s flexible regions) to firmly lock onto AP1 or AP2. And as for SH3 binding, Nef uses its proline-rich region to bind to AP1 and AP2. However, even though AP1 and AP2 binding involves also residues of the preceding flexible N-terminal arm, Nef binds these targets in very different ways. In case of AP1, Nef binds to the μ1 subunit, and part of the MHC-I cytoplasmic tail is sequestered in a groove formed between AP1 and Nef. Conversely, Nef binds to the AP2 α2, β2 and σ2 subunits. Moreover, Nef serves as an adaptor between CD4 and AP2, by using its E_160_xxxLL motif, located in the Nef core domain loop, as mimicry of acidic dileucine motifs of cellular AP2 cargos, while using a core-domain binding pocket to engage part of the CD4 cytoplasmic tail (34,37). However, despite these differences, the binding sites of CD4 and MHC-I on Nef partly overlap (37).

Collectively, these studies show that Nef’s strong and polyvalent interactions result from the combination of (i) the multivalency intrinsic to flexible protein-protein interaction motifs and (ii) the increased specificity and affinity (due to an increased binding surface) of a 3D folded domain. Nef’s composition of 50% tertiary structure and 50% flexible regions constitutes an optimal structural framework for this strategy. However, the resulting high number of cellular targets of Nef, many of which remain structurally uncharacterised, raises the question if binding to one target affects the associations of Nef with other targets. More specifically, does Nef have evolved allosteric mechanisms for creating synergy between sets of ligands that contribute to the same cellular function while excluding others? Here, we combined structural and functional studies to identify cross-reactivity in selected intra- or intermolecular associations.

## Results and discussion

### The flexible regions of Nef have specific thermodynamic effects on SH3 binding

We designed several HIV-1 Nef constructs to assess the influence of the flexible regions for binding to SH3 domains. In addition to full-length LAI Nef, we prepared (i) a deletion of the N-terminal arm that still retained the W_57_L motif that was found to be implicated in CD4 binding and that was seen in contact with the Nef hydrophobic pocket in an NMR study (19) (residues 56-205; WLΔN), (ii) a deletion of the N-terminal that lost the W_57_ and only retained L58 (residues 58-205; LΔN), (iii) a deletion of the N-terminal arm that has lost the WL motif (residues 60-205; ΔN), and (iv) a ΔN construct that had additionally a truncation of the core loop to eliminate its E_160_xxxLL endocytosis motif (residues 60-158;174-205; ΔNΔL), (**Fig 1A**). We also included full-length SF2 Nef to test the isolate-specificity of the effects. Compared to the LAI isolate, SF2 Nef has a four amino acid insertion in the N-terminal region and a T71R substitution that was shown to non-specifically enhance binding to Src family SH3 domains (28,38) (**Fig 1B**).

To test, in addition, the effect of the SH3 ‘R’ residues on Nef binding, we included SH3 mutants that were engineered to have different side chains at this position. We used Fyn SH3 wild-type (where the ‘R’ position is a long and positively charged arginine, Fyn_R96_), and an Hck-like mutant where this position is substituted by a medium-sized hydrophobic isoleucine (Fyn_R96I_), and a *Rous sarcoma* virus v-Src like large and hydrophobic tryptophan (Fyn_R96W_) (**Fig 1C**).

Using isothermal titration calorimetry (ITC), we measured the thermodynamic parameters of the three SH3 variants to different Nef constructs (**Table 1**). As expected, our ITC data showed a 1:1 binding event with the hydrophobic substitutes Fyn_R96I_ and Fyn_R96W_ displaying markedly better affinities than Fyn_R96_. However, our titration series provided additional insights:

i. Consistently across the LAI titrations, the thermodynamics of binding of Fyn_R96I_ were similar to those of Fyn_R96W_, with however Fyn_R96W_ having a more favourable enthalpy (ΔH) and less favourable entropy (ΔS) contributions. Hydrophobic interactions are normally providing a positive contribution to ΔS. Hence, the generally less favourable ΔS contribution of the most hydrophobic ‘R’ mutant Fyn_R96W_ was unexpected and suggested that additional factors influenced the binding.
ii. Truncated Nef constructs bound ~fourfold stronger to Fyn_R96W_ than to Fyn_R96I_. Conversely, both full-length Nef, LAI and SF2, associated significantly tighter with Fyn_R96I_ than with Fyn_R96W_. SF2 Nef displayed a higher affinity for all SH3 variants than full LAI Nef, which, in turn bound Fyn_R96I_ better than its truncation variants. Due to the low affinity of the Fyn_R96_ SH3 domain, our measurements did not allow us to detect precise changes between truncated and full-length LAI, but can rule out that gross affinity changes occurred. These observations suggested that the N-terminal residues 1-59 affected the affinity of SH3 binding specifically with respect to the stereochemical nature of the ‘R’ position.
iii. Fyn_R96_ bound ~tenfold weaker to the truncated LAI constructs than Fyn_R96I_. This difference to the next best SH3 variant increased in full-length Nefs to 50-fold (LAI) and 200-fold (SF2). Although precise thermodynamic parameters could not be established for Fyn_R96_ due to the low C-values of the Fyn_R96_ binding curves, this observation suggested that the presence of the N-terminal segment augmented the level of discrimination between Fyn_R96_ and the other two variants.
iv. WLΔN produced significantly different thermodynamics than the other LAI titrations. WLΔN showed both favourable ΔH and ΔS, whereas in the other LAI titrations (containing more or less flexible regions) a ~twofold more favourable ΔH compensated for an unfavourable ΔS contribution. This anomaly suggested that the W_57_L dipeptide promotes a particular thermodynamic contribution to SH3 binding, that disappears once the full N-terminal segment is present.

**Table 1.** Influence of flexible regions on the thermodynamics of SH3 binding by Nef.

| Nef : Fyn | N | Kd (μM) | $\Delta G$ (kJ/mol) | $\Delta H$ (kJ/mol) | $T\Delta S$ (kJ/mol) |
| --- | --- | --- | --- | --- | --- |
| $\Delta N\Delta L$ : Fyn<sub>R96</sub> \$ | 1* | 24.3±15.0 | -26.3±1.50 | -14.00±10.0 | 12.33±10.0 |
| $\Delta N\Delta L$ : Fyn<sub>R96I</sub> | 0.97±0.10 | 3.86±0.60 | -31.0±0.40 | -34.0±5.67 | -3.02±5.56 |
| $\Delta N\Delta L$ : Fyn<sub>R96W</sub> | 0.98±0.04 | 0.78±0.22 | -34.9±0.76 | -46.7±3.70 | -11.8±2.94 |
| $\Delta N$ : Fyn<sub>R96</sub> \$ | 1* | 83.3±15.0 | -23.3±1.50 | -34.3±10.0 | -11.0±10.0 |
| $\Delta N$ : Fyn<sub>R96I</sub> | 0.87±0.03 | 5.06±0.96 | -30.3±0.47 | -39.7±0.34 | -9.40±0.73 |
| $\Delta N$ : Fyn<sub>R96W</sub> | 0.90±0.02 | 1.28±0.45 | -33.7±0.88 | -56.5±8.68 | -22.7±9.56 |
| WL$\Delta N$ : Fyn<sub>R96</sub> \$ | 1* | 50.8±15.0 | -24.5±1.50 | -14.2±10.0 | 10.3±10.0 |
| WL$\Delta N$ : Fyn<sub>R96I</sub> | 1.01±0.05 | 7.55±1.57 | -29.3±0.54 | -21.5±2.18 | 7.77±2.71 |
| WL$\Delta N$ : Fyn<sub>R96W</sub> | 0.94±0.04 | 1.45±0.40 | -33.4±0.76 | -29.3±0.37 | 4.16±0.52 |
| LAI : Fyn<sub>R96</sub> | 1* | 58.5±8.44 | -24.2±0.68 | -45.9±10.5 | -21.7±10.8 |
| LAI : Fyn<sub>R96I</sub> | 0.93±0.01 | 0.79±0.08 | -34.9±0.23 | -40.7±1.07 | -5.80±1.30 |
| LAI : Fyn<sub>R96W</sub> | 0.96±0.02 | 1.43±0.10 | -33.4±0.15 | -50.0±1.29 | -16.6±1.41 |
| SF2 : Fyn<sub>R96</sub> | 1* | 28.7±8.72 | -26.1±0.76 | -59.2±10.3 | -33.2±11.01 |
| SF2 : Fyn<sub>R96I</sub> | 0.98±0.04 | 0.02±0.00 | -43.5±0.07 | -77.2±5.68 | -33.7±5.75 |
| SF2 : Fyn<sub>R96W</sub> | 0.91±0.03 | 0.12±0.03 | -39.5±0.58 | -56.6±6.00 | -17.1±5.58 |

Isothermal titration calorimetry data were established at 25 °C by titrating Fyn SH3 domains from the syringe onto Nef in the measurement cell (see Methods). Values given are mean ± SD, n=3, except for the following two cases: \*, the C-value of the titration was too low to establish N, and N has been fixed to 1 based on the known Nef : SH3 stoichiometry; \$, Due to low heats and low C-values, instrument-provided errors appeared underestimating the real uncertainty of the data analysis. Given errors were estimated based on observed variations in data analysis. ITC data are shown in (S1 Fig).

### The presence of SH3 domains influences binding of the N-terminal Nef helix

Our ITC data suggested that the presence of the N-terminal 59 Nef residues can influence the association with SH3 domains. This Nef region contains an amphipathic helix (termed here H1, residues 15-22 in LAI, **Fig 1A**) that helps to anchor Nef to charged lipid headgroups (23). In some structural studies of full-length Nef, this H1 helix was observed to bind back to the hydrophobic groove formed by the core helices α1 and α2 and the connecting residues (herein, we refer to this region as the α1-α2 groove)(PDB ids 6cri, 4en2, 6cm9). This α1-α2 groove is adjacent to, but not overlapping with, the SH3 binding site (35,36).

We synthesized a Nef fragment containing H1 (residues 8-26) and tested its affinity for the various Nef constructs, in the absence or presence of the SH3 Fyn_R96I_ and Fyn_R96W_ variants. To assure that most Nef was bound to the SH3 domain, we used 50 μM (approximately 10 x *K_d_*) as a concentration for the SH3 domains. We did not include the Fyn_R96_ SH3 here, because its low affinity precluded us from reaching 10 x *K_d_* in this experiment.

Using MST, we observed a low affinity of *K_d_* ~300 μM for H1 binding to the core domain–only Nef ΔNΔL. The presence of additional flexible regions further decreased this affinity: The core-loop containing Nef ΔN bound H1 with a *K_d_* of 700 μM, Nef WLΔN bound H1 with K_d_ > 1 mM and full-length LAI and SF2 did not show binding at all. We also noted that the presence of either Fyn_R96I_ or Fyn_R96W_ SH3 domains increased the affinity for Nef ΔNΔL and ΔN ~twofold (**Fig 2A-F, S1 Table**). However, WLΔN only displayed a clear twofold increase in presence of Fyn_R96I_, and both full-length Nefs failed to show an increase in H1 binding for either of the SH3 domains. Moreover, we also found that the incubation of ΔNΔL with H1 significantly increased its affinity for Fyn_R96I_ (**Fig 2G, S1 Table**).

**Fig 2.**
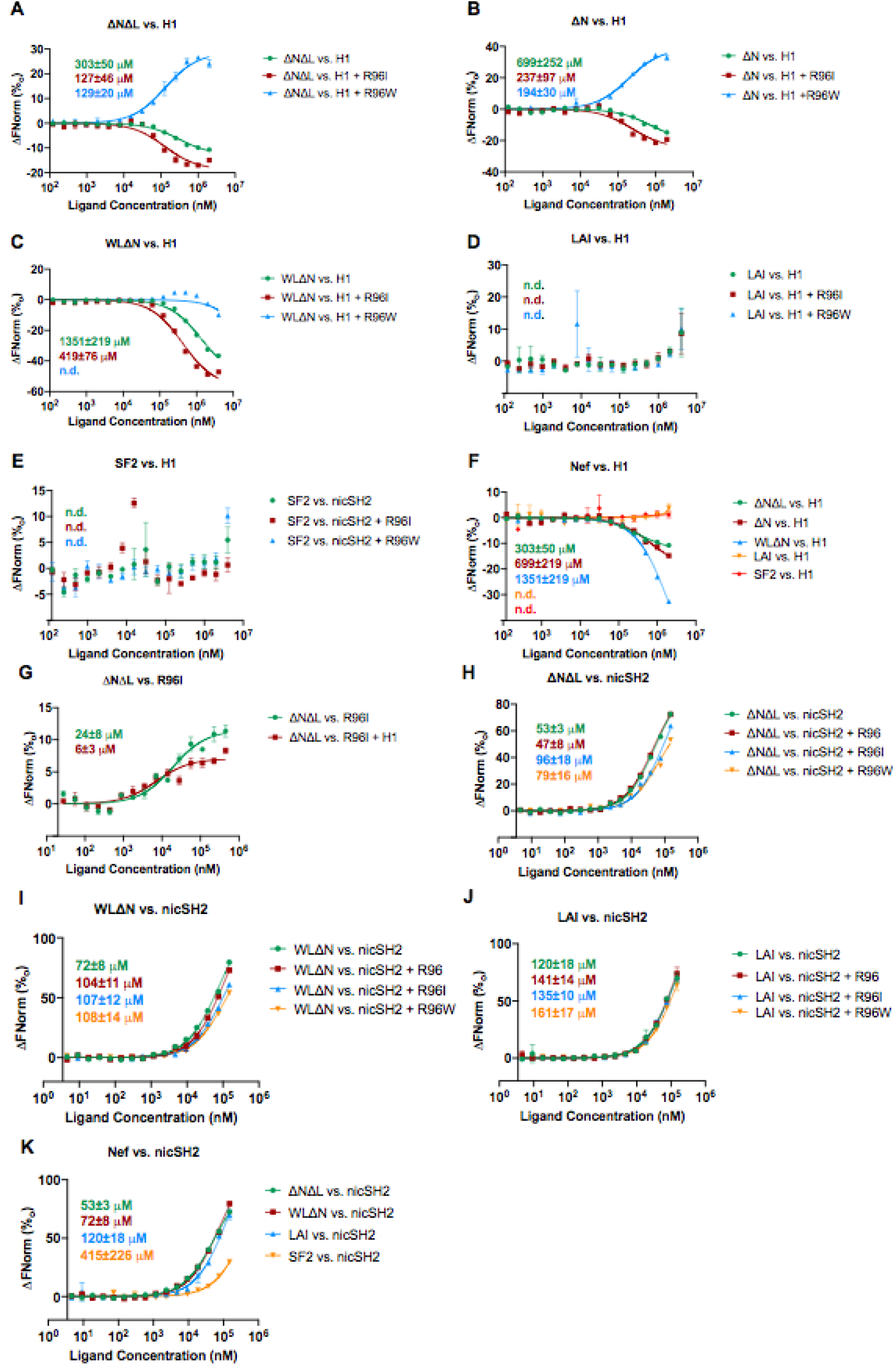
Influence of Nef’s flexible regions on inter- and intramolecular interactions. MST data were collected at room temperature. Nef was labeled and used at a fixed concentration of 10-50 nM. 50 μM SH3 domains were mixed with labelled Nef before adding the H1 or nicSH2.

Collectively, these data demonstrated that the presence of SH3 domains can enhance the association of H1 with the hydrophobic α1-α2 groove next to the SH3 binding site and *vice versa*. However, the presence of flexible Nef regions decreased binding of H1. The absence of H1 binding to full-length Nefs can be explained by the H1 sequence contained in the full-length Nefs out-competing the H1 peptide. Moreover, the W_57_L motif was found back-binding to the α1-α2 groove in the apo Nef NMR structure (2nef; (19)), explaining its competition with H1. However, competition with the core loop alone was unexpected and suggested that the core loop also loosely associates with the same Nef region.

A previous NMR study observed that the presence of the Hck SH3 domain (with an isoleucine in the ‘R’ position) enhanced the affinity of Nef for a peptide encompassing the CD4 di-leucine motif by a factor of two (from *K_d_* ~1 mM to ~ 500 μM) (39). We attempted to reproduce these data using MST and our Nef constructs (with or without SH3 domains). However, the CD4 affinities were below the detection limit of our MST assay in all cases (**S2 Fig**).

To probe whether this non-specific synergy also existed between SH3 domains and other Nef ligands, we tested the association between Nef and the C-terminal fragment of the p85 subunit of the PI3 kinase. The C-terminal region encompassing the inter-SH2 coiled-coil domain (iSH2) was previously shown to bind to HIV-1 SF2 Nef in cellular studies (40–42). However, the structural basis of this association is currently unknown. We observed binding of our LAI and SF2 Nef constructs to the p85 nSH2-iSH2-cSH2 fragment (nicSH2) with *K_d_s* in the 10s - 100 μM range (**Fig 2H-K, S1 Table**). We noted a trend of slightly decreasing affinity with the increasing length of the flexible regions, and of a slightly negative impact of the presence of R96I/W SH3 domains on the nicSH2 affinity. Although the low affinities and hence measurement uncertainties precluded us from obtaining precise affinities, we could conclude that the presence of the SH3 domains did only have neglectable effects in our assay and did not notably increase binding between Nef and p85 nicSH2.

In summary, our data established that SH3 binding to Nef (in particular Fyn_R96I_) enhanced the association of H1 to the hydrophobic α1-α2 groove of Nef. In turn, binding of H1 to the α1-α2 groove also enhanced that affinity of an SH3 domain to Nef. The α1-α2 groove is located next to the SH3 binding site but does not overlap with it. Hence, binding of one ligand to one site appears to allosterically stabilise the binding site of the other ligand. Such a stabilisation could be, for example, through paying the entropic penalty of going from the free to bond state. However, SH3 binding did not enhance the association of at least one other ligand, the p85 C-terminal region, showing that the affinity-enhancing effect of SH3 domains does not cover all Nef associations. Further, our observation that the Nef core loop and W_57_L motif can compete with the H1 motif suggested that these flexible regions can loosely associate with the Nef hydrophobic groove in absence of the H1 helix.

### Establishing the structural data for the binding features of SH3 domain variants

To identify the structural basis underlying the observed Nef : SH3 binding characteristics, we wanted to compare apo and Nef-bound structures for all three Fyn SH3 domains, Fyn_R96_, Fyn_R96I_, and Fyn_R96W_, as well as for Hck SH3. Available crystal structures used either an equivalent of the WLΔN Nef construct (an N-terminal truncation retaining the W_57_L motif), or the LΔN Nef construct which is missing W_57_ but contains L58. The following crystal structures were available: WLΔN:Fyn_R96I_ (1EFN (20)), LΔN:Fyn_R96_ (PDB id 1AVZ (22)), LΔN bound to the Hck SH2-SH3 fragment (4U5W, (24)), and the apo structures of Fyn_R96_ (1SHF, (43)) and Hck SH3 (1BU1, (27)). To obtain the missing structural data, we determined seven additional crystal structures, namely full-length SF2 Nef bound to Fyn_R96I_, of LΔN bound to Fyn_R96W_ and several structures of unliganded Fyn_R96I_ and Fyn_R96W_ (**S2-S4 Tables**).

Additionally, we ran triplicate 200 ns molecular dynamics (MD) simulations of Nef in different free and SH3-bound states (both based on the same WLΔN:Fyn_R96I_ complex taken from 1EFN) (**Supplemental Methods and Data**).

### SH3 binding decreases the dynamics of specific Nef regions

Our interaction assays showed that binding of the SH3 domain increased the affinity of Nef for H1 and vice versa, suggesting an allosteric cooperativity. To rationalize this observation, we carried out MD simulations comparing apo-WLΔN Nef (here denoted Nef-A) with Fyn_R96I_ complexed with WLΔN Nef (Nef-C). We noted that Nef-C had a lower calculated van der Waals and electrostatic energy, and showed decreased overall mobility compared to Nef-A (root mean square fluctuations, RMSF were 2.52 ± 1.88 Å for Nef-A and 2.16 ± 1.32 Å for Nef-C) (**Fig 3A-D), Supplemental Methods and Data**), indicating that the SH3 domain stabilised and rigidified Nef. The RMSF differences were greatest in the residues of the N-terminal arm and the core domain loop (**Fig 3B)**. Within the core domain, the residues 75 to 80 showed strongest stabilisation upon SH3 binding, as expected from direct interaction with the SH3 domain (**Fig 3B,E)**. This stabilisation extended to the underlying residues 111 to 126, including the C-terminal part of the α2 helix. Additionally, Nef residues 101 to 102 and residue 138 fluctuate less in the SH3 domain complex (**Fig 3F,G)**. Hence, the SH3 domain stabilised a large part of the H1 binding site and a region that interacts with CD4 (37) and contributes to Nef:Nef crystal contacts (observed in PDB 1EFN, 1AVV, 1AVZ, 4USW, 3REA, 3D8D), and possibly to dimerisation *in vitro* (44) (**Fig 3A**). Together these results suggested that SH3 binding selectively decreases the dynamics of Nef regions involved in protein-protein interactions.

**Fig 3.**
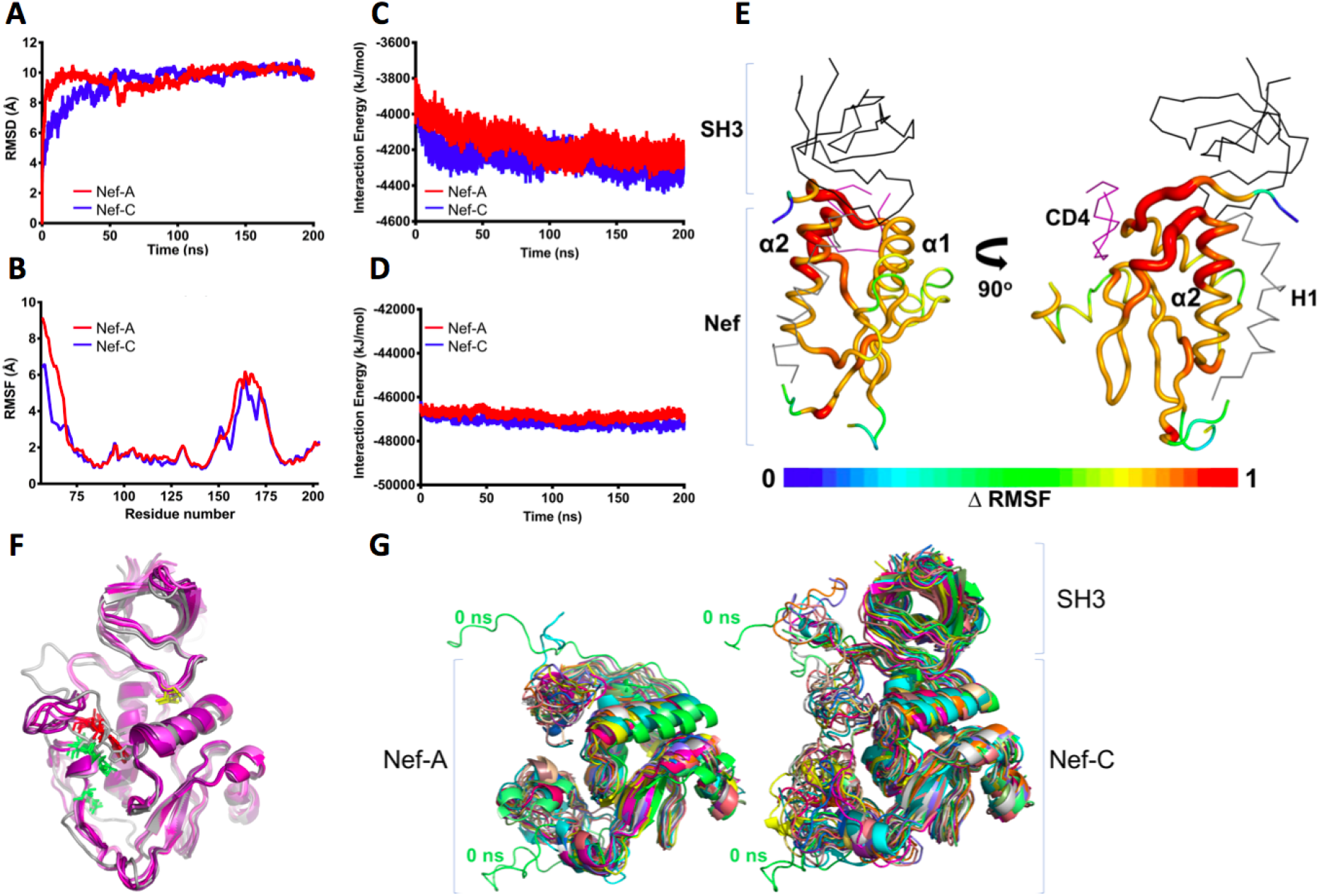
**A)** Average Cα RMSD values as a function of time for Nef-A and Nef-C. **B)** Average CαRMSF values vs residue numbers for Nef-A and Nef-C. **C)** Average Lennard-Jones and **D)** Coulombic components of the interaction energy monitored over time in Nef-A and Nef-C. **E)** B-factors showing ΔRMSF between Nef-A and Nef-C from 50-200 ns on the 3D structure. The gradient scale shows least fluctuating regions in blue and more fluctuating regions in red. SH3 domain is shown in black, CD4 is shown in magenta and H1 is shown in gray. **F)** Residual electron density in 1EFN was used to propose a speculative model for concerted back-binding of W_57_L (red) and E_160_xxxLL (green) into the a1-a2 groove. The initial model (grey) is superimposed onto structures obtained in MD simulations at 170 - 200 ns (coloured magenta to dark magenta). I96 is highlighted in yellow. **G)** (***Left***) Representative snapshots of every 10 ns for the 3D structure of Nef-apo and (***Right***) 3D structure of Nef:SH3 complex. Nef-A and Nef-C used for structural analysis using molecular dynamics are marked showing stable conformation over the course of simulation.

### Sequential associations of the flexible regions with the hydrophobic Nef α1-α2 groove

Our ITC data showed a clear difference in thermodynamic signature for SH3 domains association with Nef WLΔN as compared to the other LAI fragments. Furthermore, our MST data showed that the association of H1 is increasingly hampered by the presence of the core loop, by the W_57_L motif, and by the additional presence of the N-terminal residues 1-56. Given that H1 binds to the α1-α2 groove, the competition with H1 suggested that the core loop and the W_57_L motif also loosely associated with the α1-α2 groove.

Indeed, the hydrophobic α1-α2 groove was found occupied by intramolecular protein regions in previous structural analyses. In solution NMR studies of a truncated apo-Nef (equivalent to Nef WLΔN plus a deletion of the di-Leu endocytosis motif, as in our ΔNΔL mutant), W_57_ from the N-terminal extension was found to bind back to the α1-α2 groove (2NEF; (19)). W_57_L was also associated with the α1-α2 groove in a crystal structure of SF2 Nef bound to an engineered high-affinity SH3 domain (3REA), but in a different structural arrangement (45). In crystallographic studies of the core domains of HIV-1 and SIV Nef, the α1-α2 groove was found occupied by hydrophobic residues of a helical endocytosis motif situated in a long loop emerging from the Nef core domain of an adjacent molecule in the crystal lattice (SIV: 5NUH, 5NUI; HIV: 3RBB). However, the positions of the endocytosis motifs in HIV and SIV Nef in these crystals were different (46,47). Also, none of the core loop motifs was found associated with the Nef core domain *in cis*.

Upon reinspection of the LAI Nef_T71R_ WLΔN structure bound to Fyn_R96I_ (1EFN), we found strong and continuous residual density in the α1-α2 groove in both Fyn_R96I_-bound Nef molecules of the asymmetric unit (**S3 Fig**). In both molecules of the asymmetric unit, the electron density region closest to the position of W_57_ in the NMR model provides a good fit for a tryptophan side chain while the adjacent density fits a leucine, suggesting that this is the location of the W_57_L motif (**S3 Fig**). Assuming this position, the remaining density cannot be attributed to the residues of the N-terminal region between W_57_ and T71, and might correspond to the extremity of the core loop. Thus, we speculate that in the WL-containing Nef forms, the WL motif and the core loop might jointly occupy the α1-α2 groove, with the WL motif directly interacting with the core loop, and stabilising it. A model for such a ‘closed’ form, built based on residual electron density in 1EFN, was stable in molecular dynamics runs supporting that this conformation is stereochemically possible (**S3 Fig**).

In LAI Nef LΔN:SH3 complexes crystallized in the same space group (PDB ID 1AVZ and 4D8D, with Fyn_R96_ or Fyn_R96W_, respectively), this electron density is replaced by water molecules in both Nef of the asymmetric unit (**S3 Fig**), suggesting that the W_57_L motif was at least in part responsible for the observed unattributed electron density. A W_57_L-stabilised association of the core loop is also supported by a relative protection of this region from hydrogen exchange (48). Additionally, in our MD simulations, we noted a clear tendency of the WLΔN Nef core loop and N-terminal arm (residues 57-71) to go from their extended conformations of the initial model towards a ‘closed’ conformation where they approach the α1-α2 groove as the simulation proceeded (**Fig 3A**). Hence, our simulations suggested a tendency of these flexible residues to regroup towards the α1-α2 groove.

In combination, these observations propose a model to rationalise our ITC and MST data. In this model, the core loop alone and the W_57_L motif alone only associate very loosely and dynamically with the α1-α2 groove. When both regions are jointly available, this association becomes more pronounced. However, both are outcompeted by H1 (when tethered to the core). Further support for the capacity of H1 to outcompete other intramolecular regions comes from our survey of available experimental models (6CRI, 4EN2, 6CM9) (**S4 Table**). This survey showed that whenever the H1 motif is present in Nef, it occupies the α1-α2 groove instead of W_57_ or the core loop. In the low-resolution crystal structure of full-length LAI bound to Fyn_R96I_ that we determined the α1-α2 groove was partly occluded by adjacent molecules in the crystal lattice, further supporting that also the H1:core association is of low affinity only and can be displaced easily.

### The Fyn_R96W_ mutation alters the SH3 RT loop hydrogen-bond network

In our ITC titrations, we observed that increasing the hydrophobicity of the SH3 ‘R’ position from isoleucine to tryptophan resulted in a less favourable entropy contribution, suggesting that the incorporation of the tryptophan caused additional loss of degrees of freedom upon binding (**Table 1**). Therefore, we compared the RT loop features of our apo-Fyn_R96W_ crystal structure with those of the free states of Fyn_R96_, Fyn_R96I_ and of Hck SH3. We observed that the apo Fyn_R96_ and Fyn_R96I_ structures had a similar strong RT loop–rigidifying network of six hydrogen bonds (**Fig 4**). Conversely, the NH group of the Fyn W96 disrupts the RT loop hydrogen bond network, producing an RT-loop that is less stabilised by hydrogen bonds in the free state, and hence more flexible. Consequently, the Fyn_R96W_ variant loses more entropy upon fixation of the RT loop in the Nef interface than does Fyn_R96I_, explaining the less favourable entropy contribution to binding of Fyn_R96W_. The poorly stabilised RT loop of Fyn_R96W_ is akin to the one of Hck SH3, and Hck SH3 was previously shown to also have a less favourable enthalpy contribution to LAI Nef LΔN binding compared to other SH3 domains, including Fyn_R96_ (27). Conversely, fewer hydrogen bonds need to be broken in Fyn_R96W_ upon Nef binding, than in Fyn_R96I_ or Fyn_R96_, Fyn_R96W_, explaining the observed greater favourable enthalpy contribution of Fyn_R96W_.

**Fig 4.**
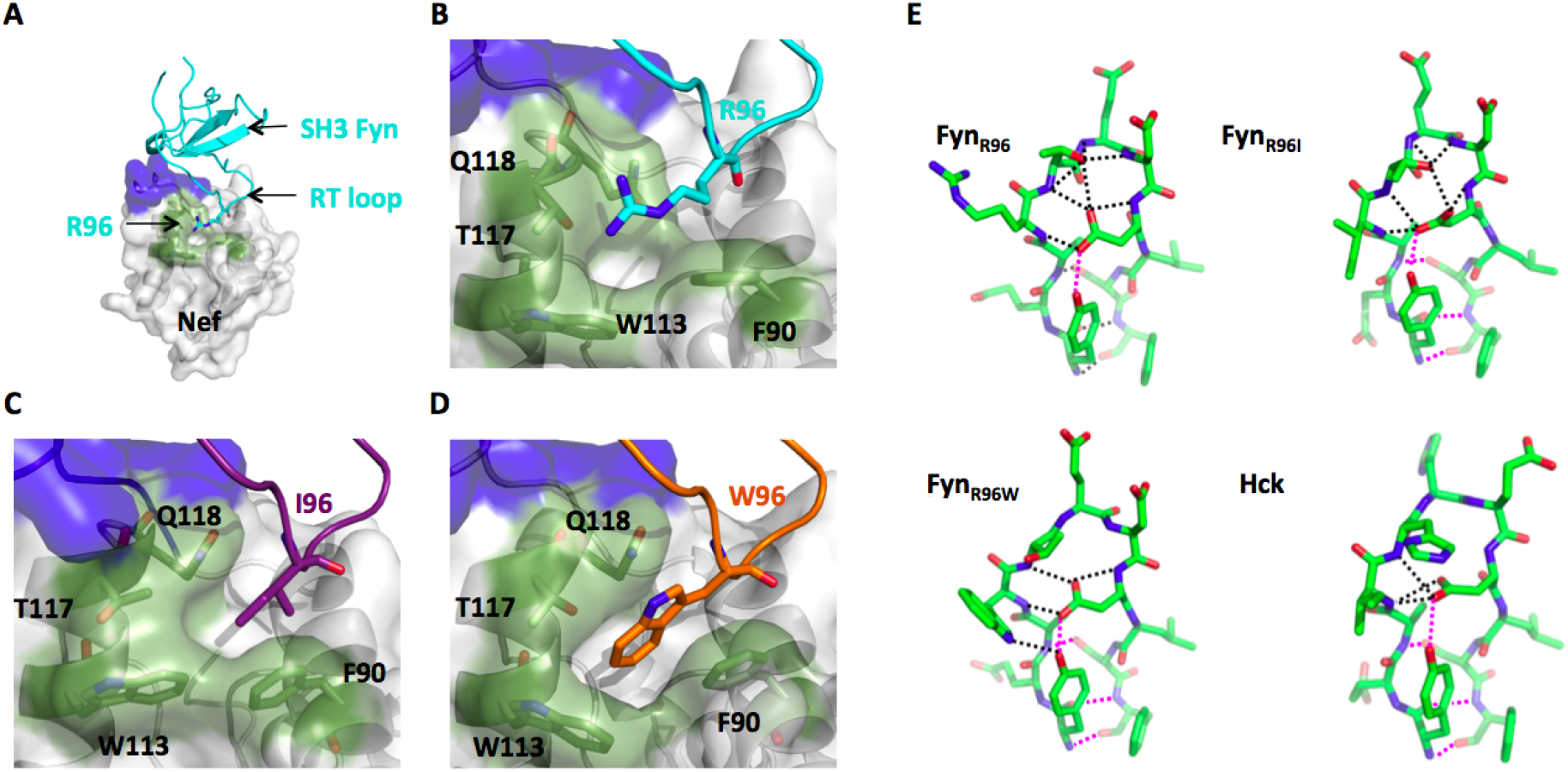
Crystal structures of the Nef core:Fyn SH3 complexes (left) and of the SH3 domain RT loop (right) are shown. The Nef P_72_xxPxR (blue) and tertiary regions (green) are shown on the Nef surface. **A)**. Global view of the LΔN:Fyn_R96_ SH3 complex (PDB 1AVZ). **B-D)** Expanded views of the tertiary interaction between SH3 residue 96 and the cognate Nef core pocket. (B) LΔN:Fyn_R96_ (1AVZ), **C)** WLΔN:Fyn_R96I_ (1EFN), and **D)** LΔN:Fyn_R96W_ (4D8D). **E)**: RT loop hydrogen bond network of unliganded SH3 domains. Hydrogen bonds are indicated by dotted lines. Those shown in dark black need to be broken upon binding to Nef core. Hydrogen bonds shown in magenta are preserved in the Nef core:SH3 complex. For Hck SH3, preserved and broken hydrogen bonds were evaluated from a computationally docked LΔN:Hck SH3 complex (22). Fyn_R96W_ SH3, but not Fyn_R96_I SH3, shows a Hck SH3-like reduced RT loop hydrogen bond network. PDB accession numbers are Hck:1BU1; Fyn_R96_:1SHF; Fyn_R96I_: 3H0I and 6IPY; Fyn_R96_W: 6IPZ.

### The Nef H1 helix selectively favours short-chain ‘R’ positions

Our binding studies suggested that the presence of the Nef N-terminal H1 helix has specific effects on SH3 domain binding: The presence of H1 increased binding of Fyn_R96I_ markedly more than Fyn_R96W_, while not having significant effects on Fyn_R96_. Our comparison of all available Nef:SH3 complexes revealed some flexibility of the relative orientation of the SH3 domains with respect to Nef, much of it can be attributed to crystallographic lattice contacts (see **Supplemental Methods and Data** and **S3 Fig**). However, the position of SH3 ‘R’-position residue relative to Nef, and the canonical P_72_xxPxR:SH3 interactions are overall well conserved (**Fig 5A**). Maintaining these key interactions is mainly possible because of the flexibility of the Nef P_72_xxPxR motif region which can buffer for positional differences compared to the rest of the Nef core. Within this context, we selected Nef:SH3 complexes that appeared unperturbed by the crystallographic lattice, and inspected the interaction between the ‘R’ position and the underlying Nef hydrophobic pocket.

**Fig 5.**
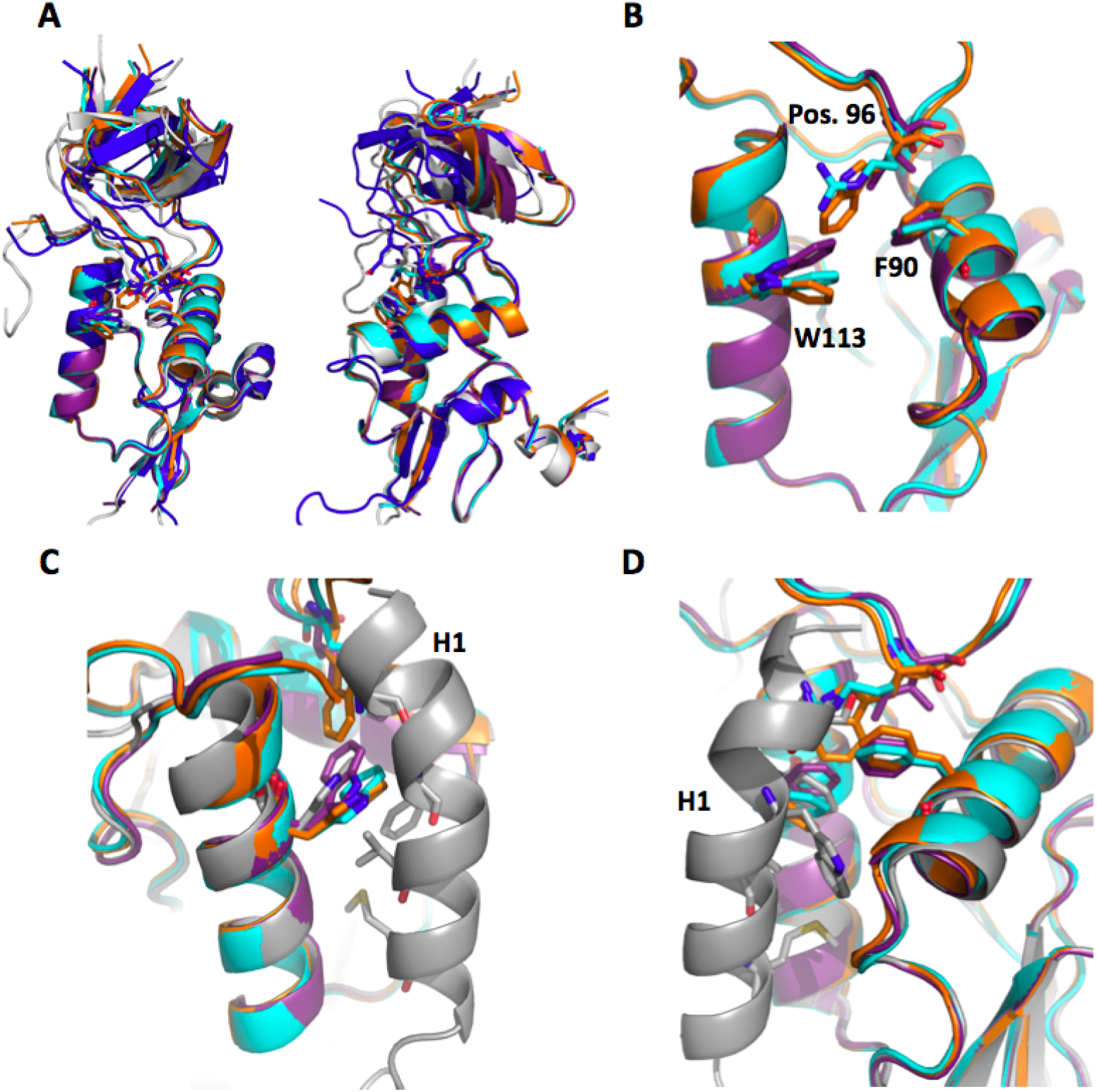
**A)** Superimposed Nef structures (1AVZ; cyan)(1efn; violet purple)(4D8D; orange)(3RBB; white)(4U5W; dark blue). **B)** Overlay of Nef LAI “gatekeeper” residues W113/F90 in relation to SH3 position 96 complexes showing Nef bound to Fyn_R96_ (1AVZ; cyan), Fyn_R96I_ (1EFN; violet purple) and Fyn_R96W_ (4D8D; orange). **C-D)** Overlap of Nef LAI “gatekeeper” residues bound to H1 (4EN2; grey).

The SH3 ‘R’ position interacts with the side chains of Nef residues W113 and F90. These side chains have been termed the ‘gatekeeper’ (47), because they separate the SH3 binding surface of Nef with the hydrophobic α1-α2 groove located between helices a1 and a2 of the Nef core domain. It is this groove that was observed binding to H1 and CD4 in previous structural studies. Hence, a change of the gatekeeper residue side-chain position as a result of their interaction with the SH3 domain will also necessarily affect the shape and size of the hydrophobic groove. Thus, the ‘R’ position of the SH3 domain could influence ligand interactions of the hydrophobic α1-α2 groove.

We noted that the side-chain position of W113, and to a lesser extent of F90, changes in response to the different SH3 orientations and to the different residues in position ‘R’. In cases where ‘R’ position is an isoleucine (Hck and Fyn_R96I_), W113 is rotated towards the SH3 domain, away from the hydrophobic crevice, as compared to apo Nef (**Fig 5B**). Conversely, in Fyn_R96W_ structures, W113 is pushed into the hydrophobic groove, whereas W113 adopts an ‘apo’ side-chain orientation in the FynR96 complex. Thus, Binding to SH3 domains with an isoleucine in position ‘R’ slightly increases the adjacent α1-α2 groove, whereas a tryptophan decreases the groove and an arginine leaves it relatively unchanged.

We then superimposed the available experimental structures of HIV-1 Nef bound to its N-terminal H1 (EM: PDB ids 6cm9, 6cri; X-ray diffraction: 4en2) with the Nef:SH3 domain complexes. In the Nef:H1 structures, the N-terminal part of H1 loosely invades the Nef region that is occupied by the ‘R’ position in the Nef:SH3 complexes, clashing much more with ‘R’ positions being a tryptophan or arginine than with an isoleucine. The more stably helical rest of H1 fills the α1-α2 groove. In this position, especially V16 and W13 of H1 are in close contact with W113 (**Fig 5C,D**). Again, an inward oriented Nef W113 (as observed in complexes with Fyn_R96I_ and Hck) would leave sufficient space for H1, while the outward pointing W113 (as seen in Fyn_R96W_ complexes) would produce (mild) clashes. From these analyses we conclude that the selective preference of an isoleucine in the ‘R’ position over an arginine or tryptophan can be explained by the direct and indirect (through the gatekeeper) impact of the H1 backbinding to the Nef core. Given that these clashes are minor, we propose that the overall affinity-enhancing effect of the N-terminal extension results from its (general) Nef-stabilising effect that is convoluted with the (specific) affinity-modulating effect on the ‘R’ position of the SH3 domain. This model can explain why full-length Nefs discriminate stronger between Fyn_R96_ and the other two SH3 variants, and reach their highest affinity for Fyn_R96I_. The additional difference between the SH3 affinities of LAI and SF2 Nef is expected to arise from the T71R substitution in SF2 Nef, that adds an additional hydrogen bond to the SH3 interaction. Indeed, in our SF2 Nef:Fyn_R96I_ crystal structure, did not show clear evidence for additional interactions between the flexible regions and the SH3 domain. However, the H1 region of SF2 Nef is characterised by the duplication of the RAEP motif situated at the C-terminus of H1, and even though this motif is not predicted to contribute to the intramolecular association, we cannot rule out that it has subtle effects on the interaction.

### Assessing the prevalence of Nef-like SH3 selectivity based on Fyn_R96_

Together with previous work, our analysis of the ‘tertiary’ association between an SH3 domain and Nef demonstrated that the combination between a linear interaction motif and the folded core domain can create opportunities for synergy and allostery, in addition to enhancing affinity and selectivity. Therefore, we wanted to assess the occurrence of a Nef-like binding mode within cellular proteins. As a high-throughput compatible proxy for this binding mode, we chose to select cellular proteins that were able to distinguish between an isoleucine and an arginine in the ‘R’ position of Src family SH3 domains. Hence, we used the Hck SH3, Fyn_R96_ SH3, and Fyn_R96I_ SH3 domains as bait for a yeast two hybrid (Y2H) analysis. First, co-transformation of cells with Nef resulted in β-galactosidase production and yeast colony outgrowth on medium lacking uracil for Hck and Fyn_R96I_ as baits but not for Fyn_R96_ (**Fig 6A**). These results were confirmed using the LacZ and HIS3 reporters for Nef:Hck and Nef:Fyn_R96_ (because Fyn_R96I_ alone was able to transactivate the lacz and HIS3 reporters, Fyn_R96I_ could not be used with these reporter systems). In contrast to Nef, the Src-associated in mitosis 68 kDa (SAM68) protein interacted with comparable strength with every SH3 domain (**Fig 6A**), indicating that the architecture of the SAM68 binding site cannot produce a Nef-like SH3 selection of the RT-loop amino acid position 96.

**Fig 6.**
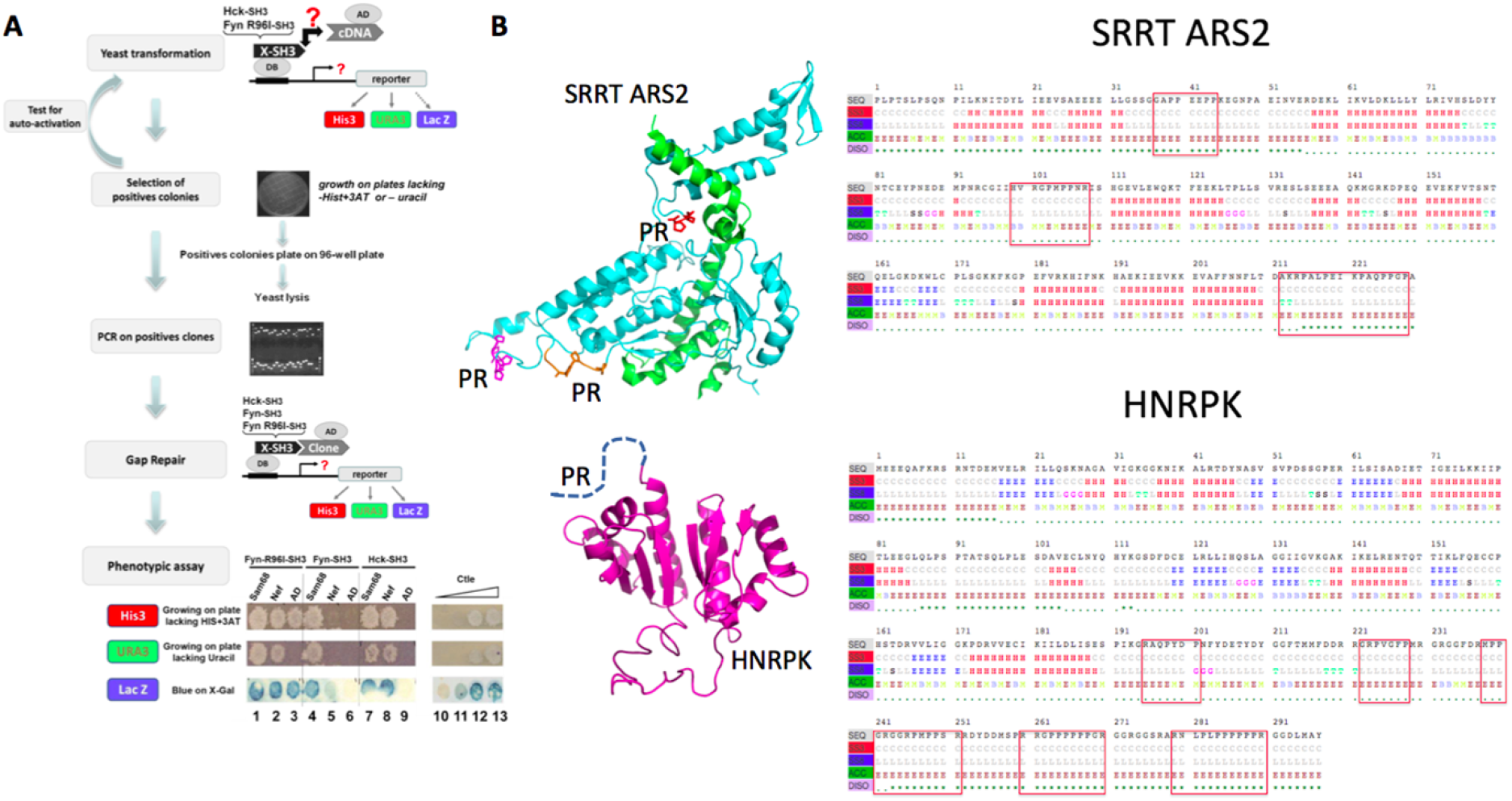
**A)** Flow chart of the protocol used to identify proteins displaying differential interaction with Hck-SH3, Fyn_R96_-SH3, and Fyn_R96I_-SH3 domains. Y2H screens were performed using either Hck-SH3 or Fyn_R96I_-SH3 domains. Positive clones were then tested against each of the Hck-SH3, Fyn(WT)-SH3, and Fyn_R96I_-SH3 domains after gap repair was performed. Binding between the two protein partners was determined by β-galactosidase activity staining (lacz) and the density of yeast colony outgrowth on medium lacking histidine (HIS3) or uracil (URA3). The strength of the corresponding interactions was evaluated by comparison with signals yielded by various known interactions (lanes 10–13) as described in Materials and Methods in the Supporting Information. Clones encoding for the HIV-1 Nef and SAM68 proteins were included as controls for differential (Nef; lanes 2, 5, and 8) or similar (SAM68; lanes 1, 4, and 7) binding to the different SH3 domains. The empty plasmid AD encoding Gal4 was used as negative control (lanes 3, 6, and 9). Of note, because the Fyn_R96I_-SH3 construct was able to transactivate the lacz and HIS3 reporters, but not URA3, only the URA3 marker was used to evaluate Fyn_R96I_-SH3-mediated interactions. **B)** *Left panel*: Homology models of SRRT and HNRPK, showing the predicted proline-rich regions (PR) that may bind to SH3 domains. *Right panel*: Prediction of secondary structure, disorder, solvent exposure and presence of SH3 binding PR motis (boxed).

To identify selective cellular binding partners for the Hck, FynR96, and Fyn_R96I_ SH3 domains, we performed Y2H screens using a human fetal library (**Fig 6A**). Interactions between these SH3 domains and cloned cDNA fragments were assessed by expression of the URA3 selection/reporter gene. Because sometimes a bait can spontaneously become a self-activator, all positive clones isolated were retested by gap repair (**Fig 6** and **Materials and Methods**). We used the gap repair step to screen for the ability of each clone to interact with each SH3 domain. To test for a Nef-like differential association, we performed two coupled serial screens. An initial screen, using the human Hck or Fyn_R96I_ SH3 domain as a bait, yielded, after gap repair, 422 and 100 clones, respectively (**S5,6 Tables**). The corresponding clones were included in a secondary screen by gap repair against the Hck, Fyn_R96_, and Fyn_R96I_ SH3 constructs, and clones displaying differential strength of β-galactosidase staining and yeast colony outgrowth on selective medium were selected and sequenced.

We found that only ten clones (9 for the screen using Hck and 3 for the Fyn_R96I_ SH3 screen, representing 2% and 3% of total clones, respectively) displayed substantially different affinities when we compared Hck and Fyn_R96I_ SH3–mediated interactions with Fyn_R96_-mediated interactions (**Table 2**). We then used bioinformatic analysis to identify putative SH3 binding motifs in these sequences (using the ELM database) and to predict the fragment’s secondary structure, solvent exposure and structural disorder (**S4 Fig**).

**Table 2.** Clones displaying differential interaction with Hck SH3 (I), Fyn SH3_R96_ (R96) and Fyn_R96I_ SH3 (R96I).

| Clones | Gene ID | Amino acids | Hck-SH3 | Fyn<sub>R96I</sub>-SH3 | PxxP | Src binding | References |
| --- | --- | --- | --- | --- | --- | --- | --- |
| **Abl-interactor 1** | 10006 | 1-282 | x |  | x | x | (51–53) |
| **Alix/PDCD6IP** | 10015 | 456-868 | x | x | x | x | (49,50) |
| **GFAP** | 2670 | 1-308 | x |  | x | x |  |
| **SRRT/ARS2** | 51593 | 546-774 | x |  | x |  | (54) |
| **HNRPK** | 3190 | 27-323 | x |  | x | x | (55) |
| **CTAGE5** | 4253 | 81-274 | x | x |  |  |  |
| **Ena/VASP-like** | 51466 | 144-278 | x |  | x | x | (56,57) |
| **FAM59A** | 64762 | 485-872 | x |  | x |  | (58) |
| **RTN3** | 10313 | 1-266 | x |  | x |  |  |
| **SMN1** | 6606 | 199-294 |  | x | x |  | (59) |

According to our RaptorX and ELM bioinformatic analysis: proteins that contained a proline-rich motif in proximity of a folded helical coiled-coil domain separated by a 20-70 amino acid linker were highlighted in light blue, proline-rich motifs close to a globular folded protein core were highlighted in light yellow, and proteins that were entirely disordered were highlighted in light red.

According to our bioinformatic analysis 5/10 clones were predicted to be almost entirely disordered, and hence cannot produce a tertiary binding site. Two clones showed a folded helical coiled-coil domain that was followed by a flexible region that contained the proline-rich motifs (ABI1 and ALIX). However, the proline-rich motifs were separated by 70 residues (ABI1) or 20 residues (ALIX (49)) from the folded part. We previously reported that ALIX achieves its capability to distinguish between an arginine and isoleucine in the SH3 ‘R’ positions through a simple ‘linear’ rather than 3D motif (49,50). Consequently, it is likely that ABI also achieves this selectivity through a ‘linear’ mechanism. GFAP is another example of a proline-rich region attached to the beginning of a (probably homotetrameric) coiled-coil structure, and might also achieve its selectivity in a linear ‘ALIX-like’ manner. Finally, only two clones (ARS2 and HNRPK) harbour PR motifs within or close to a globular folded protein core. Hence, they might offer a 3D environment that would, at least in principle, be capable of a tertiary Nef-like SH3 recognition (**Fig 6B**). Collectively, these results strongly support that only a few cellular ligands can select Src family SH3 domains according to the same RT-loop residues as does HIV-1 Nef. Yet, in most of these cases, this specificity was likely to be achieved by a linear binding mode.

## Conclusion

Despite lacking catalytic activity, SIV and HIV Nef molecules have a remarkably large functional range. And just as remarkable is the structural malleability of the flexible regions of Nef, which make up about 50% of the Nef sequence. Here, we investigated whether the combination of interactions established jointly by Nef’s folded core and flexible regions create synergy and allostery in ligand binding.

Based on our biophysical and structural analysis we propose that the flexible regions can adopt at least three different conformational states (state I - III; **Fig 7**). If the N-terminal helix H1 is available, then W_57_L and the E_160_xxxLL endocytosis motif are exposed (state I). If H1 is engaged (for example with the membrane), then the combined interaction of W_57_ and E_160_xxxLL with the core conceals these motifs (state II). If H1 is not available, and W_57_L is engaged or cleaved, then the E_160_xxxLL motif only weakly associates with the core (state III).

**Fig 7.**
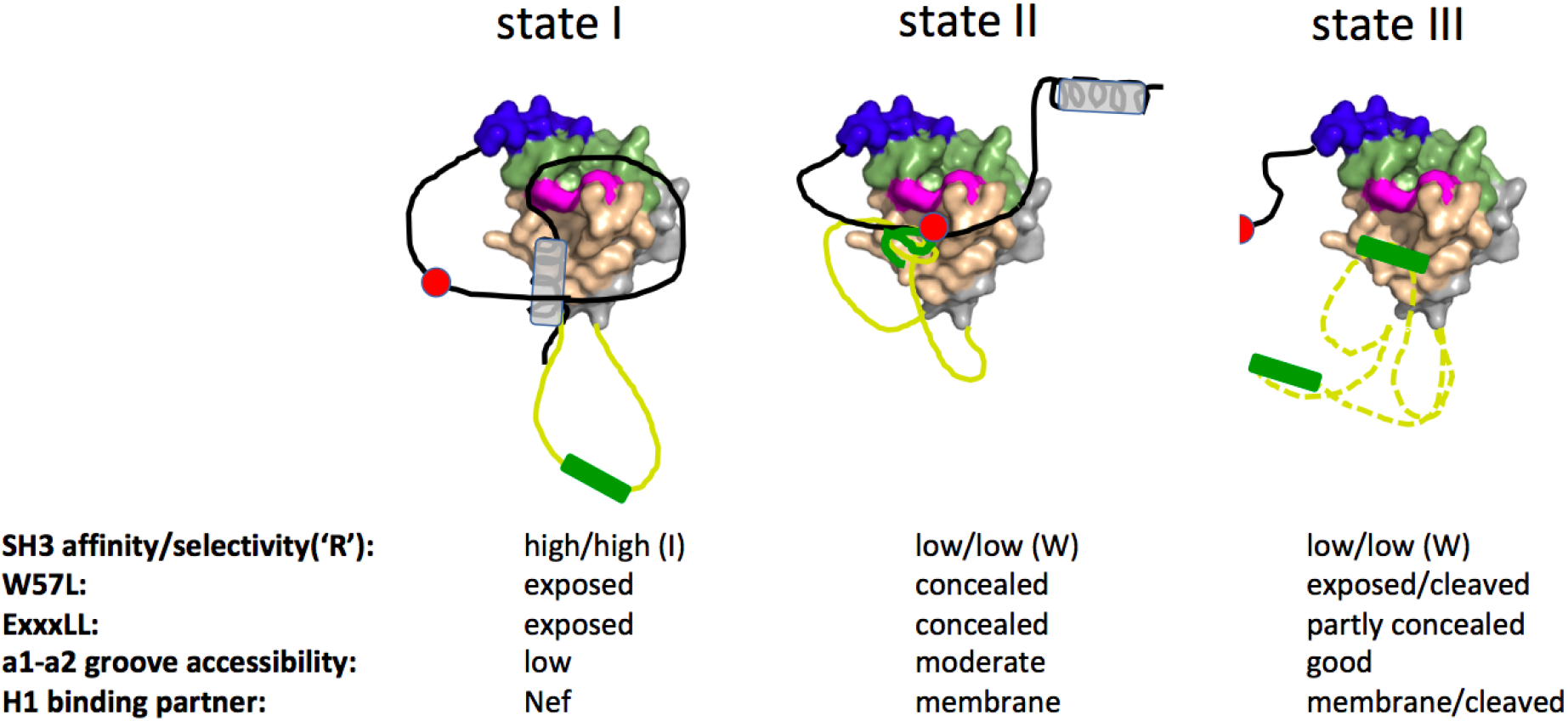
Blue: P_72_xxPxR motif; magenta: the gatekeeper residues W113 and F90; green: residues that are engaged in the tertiary SH3 interaction in addition to the gatekeeper residues. pale brown: α1-α2 groove. The remaining regions of Nef are coloured in grey. black line: N-terminal flexible region. grey helix: H1; yellow line: Nef core loop, containing the E_160_xxxLL motif (green); red sphere: W_57_L motif. The halved red sphere (state III) indicates the presence of only L58 after protease cleavage.

We further demonstrated that there is a cross-talk between the conformation of the flexible regions in the different states and the binding of some, but not all, Nef ligands. We showed that binding of SH3 domains to the tertiary Nef site affects binding of H1 to the α1-α2 groove and *vice versa*. Our analysis supports that this influence results from the combination of a non-specific dynamic coupling (binding of one ligand stabilises the Nef surface for the other ligand), and a steric interference that is specific for the SH3 ‘R’ position. This specific interference results from the size of the ‘R’ position side chain, and its interaction with the Nef gatekeeper residues. Hence, state I bound most tightly to the SH3 domain that had a mid-sized hydrophobic isoleucine in the ‘R’ position (such as Hck, Lyn and possibly Blk [where ‘R’ is a methionine]), discriminating most strongly against SH3 domains with an arginine in this position (case of Fyn, Src, Fgr and Yes). State II and III preferred the SH3 domain with a bulky hydrophobic tryptophan in the ‘R’ position (as in v-Src), but bound it less tightly and hence with less selectivity compared to wild-type Fyn. Our analysis also corroborated and extended previous findings that the affinity of SH3 domains towards Nef is additionally influenced by the extent of the hydrogen bond network that stabilises the RT loop in the apo-SH3 (27). In particular, we showed that the incorporation of the tryptophan in Fyn_R96W_ led to a loss of RT loop hydrogen bonds making it react like Hck. We also showed that state I shielded the Nef α1-α2 groove most strongly from other ligands to this site, however the states and the presence of SH3 domains did not have a measurable effect on the binding of the p85 C-terminal fragment to Nef. Our MD simulations explain this allostery, at least in part, by showing that the tertiary binding of SH3 domains stabilises specific interaction surfaces of Nef, including the one used for H1 binding. Thus, our analysis showed that the combination of a tertiary SH3 binding site with large flexible regions can provide an additional level of regulation of ligand binding. *In vivo*, Nef is myristoylated on its N-terminal glycine Nef (21,60). The presence of this fatty acid chain might further increase the number of conformational states through a myristoyl-switch mechanism and/or contribute to stabilising state I. The cross-talk between the conformational states of Nef and its ligands might help synergising certain subsets of ligands while excluding others, with potential functional implications for viral replication.

Embedded in the available functional and mechanistic data available for Nef, our findings can be extended to make the following predictions and speculations: Due to the mutual reinforcement between state I and Fyn_R96I_ SH3 domains, this state might be more predominantly found associated with the Nef function of activating Hck and Lyn. State II might present a membrane-associated and partly auto-inhibited form, in which concealing the W_57_L and the endocytosis motif prevents premature down-regulation in absence of CD4 or other surface receptors (e.g. CD3). Once specific ligands bind to the α1-α2 groove, W_57_L and the endocytosis motif would get exposed to mediate down-regulation in presence of cargo. State III would represent the conformation obtained after proteolytic cleavage by the viral protease between W_57_ and L58 (61). Cleaved Nef is predominantly found inside the HIV-1 virions. Hence, state III might facilitate tighter packing of Nef molecules within the virion, but also lead to a Nef molecule being released in infected cells that more promiscuously binds to SH3 domains and other ligands to the poorly concealed α1-α2 groove.

Our findings also raise the possibility of additional subtle allosteric mechanisms. We noted that the SH3 position ‘R’ affects the Nef W113 position, which, in turn, might affect other ligands. For example, SH2-SH3 fragments from Fyn or v-Src would not be able to form the same dimeric complex with Nef than does Hck SH2-SH3, because of clashes between Fyn R96 and W113, and/or R96 and Fyn E93 (which is required to bind to R106 of an adjacent Nef molecule in the dimer). This effect might contribute to the differential outcome between Nef interactions with either Hck or other Src kinases (24,31). Similarly, the ‘R’ position of a bound SH3 domain might favour or disfavour specific ligands to the α1-α2 groove. An additional allosteric regulation mechanism might arise from the stabilising effect of the SH3 domain on selected Nef regions. Thus, SH3 binding could promote certain Nef interactions, including some of its self-associations. SH3 binding also stabilises the Nef region involved in CD4 binding, however, the available structural models for this interaction show the proline-rich motif of Nef in a conformation incompatible with SH3 binding (37). Hence, future research needs to establish which ligands, if any, can bind Nef simultaneously with Src kinases or other SH3-domain containing ligands, such as Tec family kinases (16).

Given the versatility and potency of the combination of flexible and folded regions in Nef’s interactions, we then searched for cellular proteins that might display the same features, as illustrated by the capacity to select SH3 domains. However, our experiments revealed that only very few cellular SH3-binding proteins distinguish SH3 domains according to the same RT-loop residue as does Nef. Moreover, our bioinformatic analysis and the experimentally established example of ALIX show that the RT-loop selectivity of these cellular proteins does not necessarily require a Nef-like tertiary interaction but can be obtained through a linear peptide-like binding mode. Thus, our study infers that a Nef-like tertiary (and possibly allosterically active) SH3 binding mode is not common among cellular proteins. It is possible that Nef’s affinity/selectivity characteristics are intrinsically unsuitable for mediating SH3 binding in cellular signalling. Indeed, the tertiary recognition mode of Nef results in strong SH3 binding which not only activates Src kinases that contain a favoured isoleucine at position 96 (such as Hck or Lyn) but also perturbs the function of those with less favourable residues (such as Lck, with a serine, or Fyn and Src, with an arginine) (30,31,62,63). The Nef:SH3 domain binding mode may therefore be too strong, leading to unfavourable perturbation of several Src kinases, to be widely used in cells. The uniqueness of the tertiary SH3 binding site of Nef supports it as a selective antiviral drug target.

## Materials and Methods

### Protein production

#### Expression of protein constructs

All Fyn SH3 (Fyn_R96_, Fyn_R96I_, and Fyn_R96W_) and Nef (SF2, LAI, ΔN, ΔNΔL, and WLΔN) constructs were expressed in *Escherichia coli* Bl21 (DE3) cells with a pGEX-2T expression vector that contains an N-terminal GST tag (Fyn_R96W_; Nef WLΔN) or a pET42a expression vector that contains a C-terminal hexa-His tag (Fyn_R96_ and Fyn_R96I_) or a pET23d expression vector that contains a C-terminal hexa-HIS tag (SF2). The Nef ΔN, Nef ΔNΔL, and LAI were expressed with pJEx411c expression vector that contains an N-terminal GST tag. The bacterial cells were cultured at 37°C in 2xYT broth that contains 100 μg/ml ampicillin (all Fyn and Nef WLΔN and SF2) or 50 μg/ml kanamycin (Nef ΔN, Nef ΔNΔL, and LAI). Protein expression was induced with 0.2 mM IPTG (Fyn constructs) or 0.5 mM IPTG (Nef constructs) when the OD280 reached 0.6-0.8 and proceeded overnight at 18°C. The cells were harvested and resuspended in 50 mM Tris (pH=8.0), 200mM NaCl, 2mM EDTA (not included in His tag purification), 1mM DTT, 1 tablet protease inhibitor (Roche), 1% triton and 0.4 mg/ml lysozyme. Cells were lysed using sonication and were pelleted by centrifugation at 87,207 x g for 45 minutes at 4°C.

#### Purification of protein constructs

All proteins were purified with HIS-affinity column (Fyn_R96_, Fyn_R96I_, and SF2) or GST-affinity column using Thrombin cleavage site (Fyn_R96W_ and WLΔN) or GST-affinity column using 3C protease cleavage site (LAI, ΔN, and ΔNΔL) following standard procedures. Proteins were further purified using mono Q anion exchange chromatography and Superdex75 16/60 (GE Healthcare) size-exclusion chromatography. Protein purity was then analyzed by SDS-PAGE gel. The buffer used was 50 mM Tris (pH 8.0), 200 mM NaCl, 2 mM EDTA (Not included for HIS column), and 1 mM DTT. The protein is dialyzed in the same buffer but with 100 mM NaCl for the Mono Q column.

#### Crystallisation and structure determination

For initial crystallization experiments of the apo Fyn_R96I_ or Fyn_R96W_ SH3 domains, the SH3 domains were dialyzed against 20 mM Tris-HCl (pH 8.0), 150 mM NaCl, and 1 mM EGTA and concentrated by ultrafiltration to 5–8 mg/ml. Crystals were grown by vapor diffusion (Fyn_R96I_: sitting drop technique at 18 °C; Fyn_R96W_: hanging drop at 4 °C) by mixing 1 μl of protein solution with 1 μl of well solution (0.2 M ammonium acetate, 0.1 M sodium citrate [pH 5.6], 1.0 M lithium sulfate for Fyn_R96I_ [crystal form I]; 0.2 M ammonium acetate, 0.1 M Tris-HCl [pH 5.6], 30% v/v MPD for Fyn_R96I_ [crystal form II]; 2 M ammonium sulfate, 2% PEG 400, 0.1 M Tris [pH 9.1] for Fyn_R96W_ SH3). 25% PEG 200 was used as a cryoprotectant for Fyn_R96I_ (crystal form I). Data were recorded at 100 K (Fyn_R96I_) or 295 K (Fyn_R96W_), at λ =1.54 Å, using a Rigaku 200 X-ray generator, confocal multilayer mirrors (Osmic), and a MAR300 image plate detector. Data were integrated, merged, and scaled using Mosflm (64) and Scala (65) (**S2 Table**).

These structures were determined by molecular replacement using Molrep (66). For Fyn_R96I_ (crystal form I) and Fyn_R96W_ SH3, Fyn_R96_ SH3 was used as a template (PDB 1SHF) (43)). Fyn_R96I_ (crystal form II) was solved using Fyn_R96I_ (form I) as a template. Structures were refined using Refmac (67) and COOT (68). PDB accession codes are Fyn_R96I_: 3H0H (crystal form I) and 3H0I (crystal form II); Fyn_R96W_: 3H0F.

In a second round of crystallisation experiments, Fyn_R96I_ with a C-terminal hexahistidine tag (see (49) for cloning) crystallized in 0.1 M Citric acid pH 3.5 and 25% w/v Polyethylene glycol 3,350. Crystals were cryo-protected with 25% glycerol. The best crystal diffracted to 1.34 Å resolutions at the SOLEIL synchrotron beamline PROXIMA 2A. Fyn_R96W_ SH3 crystallized in 0.08 M sodium acetate trihydrate pH 4.6, 1.6 M Ammonium sulfate, 20% v/v Glycerol and 50 mM 18-crown-6. Crystals diffracted at best to 1.57 Å resolution at PROXIMA 2A. The structures were determined using the CCP4 online version of MoRDa (69), followed by rebuilding through BUCCANEER (70), and manual and automated (REFMAC5, PHENIX) refinement (71,72). These structures were deposited in the PDB with the accession number 6IPZ (Fyn_R96W_ SH3) and 6IPY Fyn_R96I_ SH3)).

Crystals of the HIV-1 LΔN:Fyn_R96W_ SH3 complex were obtained by hanging drop vapor diffusion at 18–20 °C. Before crystallization, stoichiometric amounts of LΔN and Fyn_R96W_ SH3 were mixed, concentrated to obtain 0.35 mM of Nef:SH3 complex, and filtered. Samples of 1 μl of this solution were mixed with 1 μl of the reservoir buffer containing 0.25–0.3 M sodium potassium tartrate, 0.5 M bicine buffer (pH 8.4), and 1 critical micelle concentration β-D-octylglucopyranoside. Cryoprotection was achieved by rapid transfer of the crystals in a cryosolvent drop containing 28% glycerol, 0.3 M sodium potassium tartrate and 0.5 M bicine buffer (pH 8.4). The crystals were flash-cooled in liquid nitrogen and data sets were collected at 100 K at the FIP beamline at the European Synchrotron Radiation Facility (Grenoble, France) using a MAR345 image plate detector. The images were processed and scaled with Mosflm (64) and Scala (65). Phases for this crystal were obtained by molecular replacement using structures of the LΔN:Fyn_R96_ SH3 and WLΔN:Fyn_R96I_ SH3 complexes as templates (PDB 1AVZ and 1EFN). The structure was refined using Refmac (71) and Phenix (72) (**S3 Table**) and has been deposited in the PDB with accession code 4d8d.

Crystals of the SF2 Nef:Fyn_R96I_ SH3 complex were obtained by sitting drop vapour diffusion at 18–20 °C. Before crystallization, stoichiometric amounts of SF2 Nef and Fyn_R96I_ SH3 were mixed, concentrated to obtain 1 mM of Nef:SH3 complex, and filtered. Samples of 1 μl of this solution were mixed with 1 μl of the reservoir buffer containing 0.10 M calcium acetate hydrate, 0.10 M sodium acetate buffer (pH 4.5), and 10% w/v PEG 4000. Cryoprotection was achieved by rapid transfer of the crystals in a cryosolvent drop containing 28% glycerol, 0.10 M calcium acetate hydrate, 0.10 M sodium acetate buffer (pH 4.5), and 10% w/v PEG 4000. All data for SF2 Nef:Fyn_R96I_ SH3 were collected at 100K at the beamline Proxima 2A at the SOLEIL Synchrotron (France), EIGER 9M detector, respectively (proposal numbers 2016 0098, 20161236, 20170193). Data from three crystals were processed, scaled and combined using XDS as implemented in the XDSme pipeline (unmerged data are available at https://doi.org/10.25781/KAUST-8CV15). Due to anisotropic diffraction, data were merged and anisotropic resolution limits established using STARANISO (73) (**S3 Table**). The structure was solved by MoRDa (69) using the PDB entry 4d8d as molecular template (Q score 0.834). The structure was manually corrected (COOT) and refined by LORESTR (74) (Ramachandran outliers/favourite 2.53/91.14%)

#### Molecular dynamics simulations

Molecular dynamics simulations in triplicates were carried out for Nef-A and Nef-C, starting from the experimental structure (PDB ID: 1EFN). MD simulations were carried out using GROMACS 2018 (75), with AMBER14SB force field (76). Each protein was inserted into a cubic box filled with TIP3P water molecules, setting a minimum distance of 10.0 Å from the protein edges to the box sides. The system was neutralized using the solution with Na+ and Cl- ions. Energy minimization was performed, followed by equilibration using isothermal ensemble dynamics (NVT) with a velocity-rescale thermostat (77) for computing velocities and positions of atoms. Further, equilibration of 2 ns with isothermal-isobaric ensemble dynamics (NPT) was carried out on the structure. Periodic boundary conditions (PBC) were applied in all X, Y, and Z directions. Final production simulations were carried out using NPT ensemble for 200 ns in triplicates. The temperature was kept constant at 300K using a velocity-rescale thermostat (77) (τT = 0.1 ps) and 1 bar pressure was maintained using a Parrinello-Rahman barostat (78) (τP = 2.0 ps). Electrostatic interactions beyond 12 Å were evaluated by the Particle-Mesh-Ewald (PME) (79). LINear Constraint Solver algorithm (LINCS) (80) was used to constrain the bond lengths. Output trajectories were analyzed using GROMACS 2018 analysis tools and PyMol (www.pymol.org).

#### Microscale thermophoresis (MST)

All proteins were labeled with the Monolith Protein Labeling Kit RED-NHS 2nd Generation. The labeled protein concentrations ranged from 10-50 nM. The unlabeled proteins were at least 10 folds higher than the expected *K_d_*. The measurements were performed at 25-50% LED power and 40% MST power. Data was analyzed using the analysis program provided by Nanotemper Technologies. All MST experiments were performed in 50 mM Tris (pH=8.0), 200mM NaCl, 2mM EDTA (not included in HIS tag purification), 1-2mM DTT at room temperature.

#### Isothermal titration calorimetry (ITC)

All ITC experiments were performed on MicroCal PEAQ-ITC machine by malvern panalytical (19-injection standard method, 25 °C). All proteins were dialyzed and degassed in 20mM Sodium phosphate (pH=7.5), 150mM NaCl, 2mM EDTA, 1-2mM TCEP. The protein concentrations in the cell ranged from 20-75 μM and were 10 folds higher in the syringe. The measurements and data were analyzed using Analysis Software provided by ITC Origin program.

#### Yeast Two-Hybrid (Y2H) screen

Y2H screens were performed as reported previously (50).

#### Bioinformatic analysis

The Y2H bioinformatic analysis was performed using the RaptorX for property prediction/structure prediction and the ELM database for identification of SH3 binding motifs (81,82).

## Acknowledgements

We acknowledge SOLEIL for provision of synchrotron radiation facilities and we would like to thank L. Chavas, P. Legrand, S. Sirigu and P. Montaville for assistance in using beamline PROXIMA 1, G. Fox, M. Savko and B. Shepard for assistance in using beamline PROXIMA 2A. We thank M-P. Struband M-T. Augé-Sénégasfor the initial cloning of some of the Fyn_R96I_ and Fyn_R96W_ mutants. We thank M. Geyer for providing the expression construct of Nef SF2. We thank the staff of the ESRF beamline BM30 (Grenoble, France) for assistance with crystallographic data collection. We also thank the KAUST Supercomputing Laboratory (KSL) for their assistance with computational resources for molecular dynamics simulations using the IBEX cluster.

## Supporting information

## References

1. Arold ST, Baur AS. Dynamic Nef and Nef dynamics: how structure could explain the complex activities of this small HIV protein. Trends in Biochemical Sciences. 2001 Jun;26(6):356–63.

2. Kirchhoff F, Schindler M, Specht A, Arhel N, Münch J. Role of Nef in primate lentiviral immunopathogenesis. Cell Mol Life Sci. 2008 Sep;65(17):2621–36.

3. Kestler HW, Ringler DJ, Mori K, Panicali DL, Sehgal PK, Daniel MD, et al. Importance of the nef gene for maintenance of high virus loads and for development of AIDS. Cell. 1991 May 17;65(4):651–62.

4. Kirchhoff F, Greenough TC, Brettler DB, Sullivan JL, Desrosiers RC. Absence of Intact nef Sequences in a Long-Term Survivor with Nonprogressive HIV-1 Infection. New England Journal of Medicine. 1995 Jan 26;332(4):228–32.

5. Deacon NJ, Tsykin A, Solomon A, Smith K, Ludford-Menting M, Hooker DJ, et al. Genomic structure of an attenuated quasi species of HIV-1 from a blood transfusion donor and recipients. Science. 1995 Nov 10;270(5238):988–91.

6. Pereira EA, daSilva LLP. HIV-1 Nef: Taking Control of Protein Trafficking. Traffic. 2016;17(9):976–96.

7. Buffalo CZ, Iwamoto Y, Hurley JH, Ren X. How HIV Nef Proteins Hijack Membrane Traffic To Promote Infection. Pierson TC, editor. J Virol. 2019 Oct 2;93(24):e01322–19, /jvi/93/24/JVI.01322-19.atom.

8. Lama J, Ware CF. Human Immunodeficiency Virus Type 1 Nef Mediates Sustained Membrane Expression of Tumor Necrosis Factor and the Related Cytokine LIGHT on Activated T Cells. J Virol. 2000 Oct;74(20):9396–402.

9. Sol-Foulon N, Moris A, Nobile C, Boccaccio C, Engering A, Abastado J-P, et al. HIV-1 Nef-induced upregulation of DC-SIGN in dendritic cells promotes lymphocyte clustering and viral spread. Immunity. 2002 Jan;16(1):145–55.

10. Schrager JA, Marsh JW. HIV-1 Nef increases T cell activation in a stimulus-dependent manner. Proc Natl Acad Sci U S A. 1999 Jul 6;96(14):8167–72.

11. Markle TJ, Philip M, Brockman MA. HIV-1 Nef and T-cell activation: a history of contradictions. Future Virol. 2013 Apr 1;8(4).

12. Swingler S, Mann A, Jacqué J, Brichacek B, Sasseville VG, Williams K, et al. HIV-1 Nef mediates lymphocyte chemotaxis and activation by infected macrophages. Nat Med. 1999 Sep;5(9):997–103.

13. Arora VK, Molina RP, Foster JL, Blakemore JL, Chernoff J, Fredericksen BL, et al. Lentivirus Nef Specifically Activates Pak2. J Virol. 2000 Dec;74(23):11081–7.

14. Renkema GH, Manninen A, Mann DA, Harris M, Saksela K. Identification of the Nef-associated kinase as p21-activated kinase 2. Current Biology. 1999 Dec 2;9(23):1407–11.

15. Briggs SD, Sharkey M, Stevenson M, Smithgall TE. SH3-mediated Hck tyrosine kinase activation and fibroblast transformation by the Nef protein of HIV-1. J Biol Chem. 1997 Jul 18;272(29):17899–902.

16. Tarafdar S, Poe JA, Smithgall TE. The Accessory Factor Nef Links HIV-1 to Tec/Btk Kinases in an Src Homology 3 Domain-dependent Manner◆. J Biol Chem. 2014 May 30;289(22):15718–28.

17. Emert-Sedlak LA, Narute P, Shu ST, Poe JA, Shi H, Yanamala N, et al. Effector kinase coupling enables high-throughput screens for direct HIV-1 Nef antagonists with antiretroviral activity. Chem Biol. 2013 Jan 24;20(1):82–91.

18. Wolf D, Witte V, Laffert B, Blume K, Stromer E, Trapp S, et al. HIV-1 Nef associated PAK and PI3-Kinases stimulate Akt-independent Bad-phosphorylation to induce anti-apoptotic signals. Nat Med. 2001 Nov;7(11):1217–24.

19. Grzesiek S, Bax A, Hu JS, Kaufman J, Palmer I, Stahl SJ, et al. Refined solution structure and backbone dynamics of HIV-1 Nef. Protein Sci. 1997 Jun;6(6):1248–63.

20. Lee C-H, Saksela K, Mirza UA, Chait BT, Kuriyan J. Crystal Structure of the Conserved Core of HIV-1 Nef Complexed with a Src Family SH3 Domain. Cell. 1996 Jun 14;85(6):931–42.

21. Geyer M, Munte CE, Schorr J, Kellner R, Kalbitzer HR. Structure of the anchor-domain of myristoylated and non-myristoylated HIV-1 Nef protein. J Mol Biol. 1999 May 28;289(1):123–38.

22. Arold S, Franken P, Strub MP, Hoh F, Benichou S, Benarous R, et al. The crystal structure of HIV-1 Nef protein bound to the Fyn kinase SH3 domain suggests a role for this complex in altered T cell receptor signaling. Structure. 1997 Oct 15;5(10):1361–72.

23. Bentham M, Mazaleyrat S, Harris M. Role of myristoylation and N-terminal basic residues in membrane association of the human immunodeficiency virus type 1 Nef protein. Journal of General Virology,. 2006;87(3):563–71.

24. Alvarado JJ, Tarafdar S, Yeh JI, Smithgall TE. Interaction with the Src homology (SH3-SH2) region of the Src-family kinase Hck structures the HIV-1 Nef dimer for kinase activation and effector recruitment. J Biol Chem. 2014 Oct 10;289(41):28539–53.

25. Lee CH, Leung B, Lemmon MA, Zheng J, Cowburn D, Kuriyan J, et al. A single amino acid in the SH3 domain of Hck determines its high affinity and specificity in binding to HIV-1 Nef protein. The EMBO Journal. 1995 Oct 1;14(20):5006–15.

26. Collette Y, Arold S, Picard C, Janvier K, Benichou S, Benarous R, et al. HIV-2 and SIV Nef Proteins Target Different Src Family SH3 Domains than Does HIV-1 Nef because of a Triple Amino Acid Substitution. J Biol Chem. 2000 Feb 11;275(6):4171–6.

27. Arold S, O’Brien R, Franken P, Strub M-P, Hoh F, Dumas C, et al. RT Loop Flexibility Enhances the Specificity of Src Family SH3 Domains for HIV-1 Nef,. Biochemistry. 1998 Oct 1;37(42):14683–91.

28. Moarefi I, LaFevre-Bernt M, Sicheri F, Huse M, Lee CH, Kuriyan J, et al. Activation of the Src-family tyrosine kinase Hck by SH3 domain displacement. Nature. 1997 Feb 13;385(6617):650–3.

29. Lerner EC, Smithgall TE. SH3-dependent stimulation of Src-family kinase autophosphorylation without tail release from the SH2 domain in vivo. Nat Struct Biol. 2002 May;9(5):365–9.

30. Trible RP, Emert-Sedlak L, Smithgall TE. HIV-1 Nef Selectively Activates Src Family Kinases Hck, Lyn, and c-Src through Direct SH3 Domain Interaction. J Biol Chem. 2006 Sep 15;281(37):27029–38.

31. Briggs SD, Lerner EC, Smithgall TE. Affinity of Src family kinase SH3 domains for HIV Nef in vitro does not predict kinase activation by Nef in vivo. Biochemistry. 2000 Jan 25;39(3):489–95.

32. Hung C-H, Thomas L, Ruby CE, Atkins KM, Morris NP, Knight ZA, et al. HIV-1 Nef Assembles a Src Family Kinase-ZAP-70/Syk-PI3K Cascade to Downregulate Cell-Surface MHC-I. Cell Host & Microbe. 2007 Apr 19;1(2):121–33.

33. Lee J-H, Ostalecki C, Zhao Z, Kesti T, Bruns H, Simon B, et al. HIV Activates the Tyrosine Kinase Hck to Secrete ADAM Protease-Containing Extracellular Vesicles. EBioMedicine. 2018;28:151–61.

34. Ren X, Park SY, Bonifacino JS, Hurley JH. How HIV-1 Nef hijacks the AP-2 clathrin adaptor to downregulate CD4. eLife [Internet]. [cited 2020 Aug 18];3. Available from: https://www.ncbi.nlm.nih.gov/pmc/articles/PMC3901399/

35. Morris KL, Buffalo CZ, Stürzel CM, Heusinger E, Kirchhoff F, Ren X, et al. HIV-1 Nefs are cargo-sensitive AP-1 trimerization switches in tetherin downregulation. Cell. 2018 Jul 26;174(3):659–671.e14.

36. Jia X, Singh R, Homann S, Yang H, Guatelli J, Xiong Y. Structural basis of evasion of cellular adaptive immunity by HIV-1 Nef. Nat Struct Mol Biol. 2012 Jul;19(7):701–6.

37. Kwon Y, Kaake RM, Echeverria I, Suarez M, Karimian Shamsabadi M, Stoneham C, et al. Structural basis of CD4 downregulation by HIV-1 Nef. Nat Struct Mol Biol [Internet]. 2020 Jul 27 [cited 2020 Aug 18]; Available from: http://www.nature.com/articles/s41594-020-0463-z

38. Saksela K, Cheng G, Baltimore D. Proline-rich (PxxP) motifs in HIV-1 Nef bind to SH3 domains of a subset of Src kinases and are required for the enhanced growth of Nef+ viruses but not for down-regulation of CD4. EMBO J. 1995 Feb 1;14(3):484–91.

39. Grzesiek S, Stahl SJ, Wingfield PT, Bax A. The CD4 determinant for downregulation by HIV-1 Nef directly binds to Nef. Mapping of the Nef binding surface by NMR. Biochemistry. 1996 Aug 13;35(32):10256–61.

40. Linnemann T, Zheng Y-H, Mandic R, Matija Peterlin B. Interaction between Nef and Phosphatidylinositol-3-Kinase Leads to Activation of p21-Activated Kinase and Increased Production of HIV. Virology. 2002 Mar 15;294(2):246–55.

41. Graziani A, Galimi F, Medico E, Cottone E, Gramaglia D, Boccaccio C, et al. The HIV-1 Nef Protein Interferes with Phosphatidylinositol 3-Kinase Activation 1. J Biol Chem. 1996 Mar 22;271(12):6590–3.

42. Kim Y-H, Chang SH, Kwon JH, Rhee SS. HIV-1 Nef Plays an Essential Role in Two Independent Processes in CD4 Down-regulation: Dissociation of the CD4–p56lckComplex and Targeting of CD4 to Lysosomes. Virology. 1999 Apr 25;257(1):208–19.

43. Noble ME, Musacchio A, Saraste M, Courtneidge SA, Wierenga RK. Crystal structure of the SH3 domain in human Fyn; comparison of the three-dimensional structures of SH3 domains in tyrosine kinases and spectrin. EMBO J. 1993 Jul;12(7):2617–24.

44. Franken P, Arold S, Padilla A, Hoh E, Strub MP, Boyer M, et al. HIV-1 Nef protein: Purification, crystallizations, and preliminary X-ray diffraction studies. Protein Science. 1997;6(12):2681–3.

45. Breuer S, Schievink SI, Schulte A, Blankenfeldt W, Fackler OT, Geyer M. Molecular design, functional characterization and structural basis of a protein inhibitor against the HIV-1 pathogenicity factor Nef. PLoS ONE. 2011;6(5):e20033.

46. Manrique S, Sauter D, Horenkamp FA, Lülf S, Yu H, Hotter D, et al. Endocytic sorting motif interactions involved in Nef-mediated downmodulation of CD4 and CD3. Nat Commun [Internet]. 2017 Sep 5 [cited 2020 Aug 18];8. Available from: https://www.ncbi.nlm.nih.gov/pmc/articles/PMC5585231/

47. Horenkamp FA, Breuer S, Schulte A, Lülf S, Weyand M, Saksela K, et al. Conformation of the Dileucine-Based Sorting Motif in HIV-1 Nef Revealed by Intermolecular Domain Assembly. Traffic. 2011;12(7):867–77.

48. Hochrein JM, Wales TE, Lerner EC, Schiavone AP, Smithgall TE, Engen JR. Conformational Features of the Full-Length HIV and SIV Nef Proteins Determined by Mass Spectrometry. Biochemistry. 2006 Jun 1;45(25):7733–9.

49. Shi X, Opi S, Lugari A, Restouin A, Coursindel T, Parrot I, et al. Identification and biophysical assessment of the molecular recognition mechanisms between the human haemopoietic cell kinase Src homology domain 3 and ALG-2-interacting protein X. Biochem J. 2010 Oct 1;431(1):93–102.

50. Shi X, Betzi S, Lugari A, Opi S, Restouin A, Parrot I, et al. Structural recognition mechanisms between human Src homology domain 3 (SH3) and ALG-2-interacting protein X (Alix). FEBS Letters. 2012 Jun 21;586(13):1759–64.

51. Shi Y, Alin K, Goff SP. Abl-interactor-1, a novel SH3 protein binding to the carboxy-terminal portion of the Abl protein, suppresses v-abl transforming activity. Genes Dev. 1995 Nov 1;9(21):2583–97.

52. Rickles RJ, Botfield MC, Weng Z, Taylor JA, Green OM, Brugge JS, et al. Identification of Src, Fyn, Lyn, PI3K and Abl SH3 domain ligands using phage display libraries. The EMBO Journal. 1994 Dec 1;13(23):5598–604.

53. Sparks AB, Rider JE, Hoffman NG, Fowlkes DM, Quillam LA, Kay BK. Distinct ligand preferences of Src homology 3 domains from Src, Yes, Abl, Cortactin, p53bp2, PLCgamma, Crk, and Grb2. PNAS. 1996 Feb 20;93(4):1540–4.

54. Thalappilly S, Suliman M, Gayet O, Soubeyran P, Hermant A, Lecine P, et al. Identification of multi-SH3 domain-containing protein interactome in pancreatic cancer: A yeast two-hybrid approach. PROTEOMICS. 2008;8(15):3071–81.

55. Lu J, Gao F-H. Role and molecular mechanism of heterogeneous nuclear ribonucleoprotein K in tumor development and progression (Review). Biomedical Reports. 2016 Jun 1;4(6):657–63.

56. Reinhard M, Jarchau T, Walter U. Actin-based motility: stop and go with Ena/VASP proteins. Trends in Biochemical Sciences. 2001 Apr 1;26(4):243–9.

57. Kwiatkowski AV, Gertler FB, Loureiro JJ. Function and regulation of Ena/VASP proteins. Trends in Cell Biology. 2003 Jul 1;13(7):386–92.

58. Tashiro K, Tsunematsu T, Okubo H, Ohta T, Sano E, Yamauchi E, et al. GAREM, a Novel Adaptor Protein for Growth Factor Receptor-bound Protein 2, Contributes to Cellular Transformation through the Activation of Extracellular Signal-regulated Kinase Signaling. J Biol Chem. 2009 Jul 24;284(30):20206–14.

59. Kurihara N, Menaa C, Maeda H, Haile DJ, Reddy SV. Osteoclast-stimulating factor interacts with the spinal muscular atrophy gene product to stimulate osteoclast formation. J Biol Chem. 2001 Nov 2;276(44):41035–9.

60. Gerlach H, Laumann V, Martens S, Becker CFW, Goody RS, Geyer M. HIV-1 Nef membrane association depends on charge, curvature, composition and sequence. Nat Chem Biol. 2010 Jan;6(1):46–53.

61. Chen Y-L, Trono D, Camaur D. The Proteolytic Cleavage of Human Immunodeficiency Virus Type 1 Nef Does Not Correlate with Its Ability To Stimulate Virion Infectivity. Journal of Virology. 1998 Apr 1;72(4):3178–84.

62. Collette Y, Dutartre H, Benziane A, Ramos-Morales F, Benarous R, Harris M, et al. Physical and Functional Interaction of Nef with Lck HIV-1 Nef-INDUCED T-CELL SIGNALING DEFECTS. J Biol Chem. 1996 Mar 15;271(11):6333–41.

63. Greenway AL, Dutartre H, Allen K, McPhee DA, Olive D, Collette Y. Simian Immunodeficiency Virus and Human Immunodeficiency Virus Type 1 Nef Proteins Show Distinct Patterns and Mechanisms of Src Kinase Activation. Journal of Virology. 1999 Jul 1;73(7):6152–8.

64. Leslie AGW. Recent changes to the MOSFLM package for processing film and image plate data. CCP4 and ESF-EACMB Newsletter on Protein Crystallography [Internet]. 1992 [cited 2020 Sep 11]; Available from: https://ci.nii.ac.jp/naid/10020054202/

65. Collaborative Computational Project, Number 4. The CCP4 suite: programs for protein crystallography. Acta Crystallogr D Biol Crystallogr. 1994 Sep 1;50(Pt 5):760–3.

66. Vagin A, Teplyakov A. MOLREP: an Automated Program for Molecular Replacement. J Appl Cryst. 1997 Dec 1;30(6):1022–5.

67. Murshudov GN, Vagin AA, Dodson EJ. Refinement of Macromolecular Structures by the Maximum-Likelihood Method. Acta Cryst D. 1997 May 1;53(3):240–55.

68. Emsley P, Cowtan K. Coot: model-building tools for molecular graphics. Acta Cryst D. 2004 Dec 1;60(12):2126–32.

69. Vagin A, Lebedev A. MoRDa, an automatic molecular replacement pipeline [Internet]. Vol. 71, Acta Crystallographica Section A: Foundations and Advances. International Union of Crystallography; 2015 [cited 2020 Sep 11]. p. s19–s19. Available from: https://scripts.iucr.org/cgi-bin/paper?a53232

70. Cowtan K. The Buccaneer software for automated model building. 1. Tracing protein chains. Acta Cryst D. 2006 Sep 1;62(9):1002–11.

71. Murshudov GN, Skubák P, Lebedev AA, Pannu NS, Steiner RA, Nicholls RA, et al. REFMAC5 for the refinement of macromolecular crystal structures. Acta Cryst D. 2011 Apr 1;67(4):355–67.

72. DiMaio F, Echols N, Headd JJ, Terwilliger TC, Adams PD, Baker D. Improved low-resolution crystallographic refinement with Phenix and Rosetta. Nature Methods. 2013 Nov;10(11):1102–4.

73. Vonrhein C, Tickle IJ, Flensburg C, Keller P, Paciorek W, Sharff A, et al. Advances in automated data analysis and processing within autoPROC, combined with improved characterisation, mitigation and visualisation of the anisotropy of diffraction limits using STARANISO [Internet]. Vol. 74, Acta Crystallographica Section A: Foundations and Advances. International Union of Crystallography; 2018 [cited 2020 Sep 11]. p. a360–a360. Available from: https://scripts.iucr.org/cgi-bin/paper?S010876731809640X

74. Kovalevskiy O, Nicholls RA, Murshudov GN. Automated refinement of macromolecular structures at low resolution using prior information. Acta Cryst D. 2016 Oct 1;72(10):1149–61.

75. Abraham MJ, Murtola T, Schulz R, Páll S, Smith JC, Hess B, et al. GROMACS: High performance molecular simulations through multi-level parallelism from laptops to supercomputers. SoftwareX. 2015 Sep 1;1–2:19–25.

76. Maier JA, Martinez C, Kasavajhala K, Wickstrom L, Hauser KE, Simmerling C. ff14SB: Improving the Accuracy of Protein Side Chain and Backbone Parameters from ff99SB. J Chem Theory Comput. 2015 Aug 11;11(8):3696–713.

77. Bussi G, Donadio D, Parrinello M. Canonical sampling through velocity rescaling. J Chem Phys. 2007 Jan 3;126(1):014101.

78. Parrinello M, Rahman A. Polymorphic transitions in single crystals: A new molecular dynamics method. Journal of Applied Physics. 1981 Dec 1;52(12):7182–90.

79. Essmann U, Perera L, Berkowitz ML, Darden T, Lee H, Pedersen LG. A smooth particle mesh Ewald method. J Chem Phys. 1995 Nov 15;103(19):8577–93.

80. Hess B, Bekker H, Berendsen HJC, Fraaije JGEM. LINCS: A linear constraint solver for molecular simulations. Journal of Computational Chemistry. 1997;18(12):1463–72.

81. Källberg M, Wang H, Wang S, Peng J, Wang Z, Lu H, et al. Template-based protein structure modeling using the RaptorX web server. Nature Protocols. 2012 Aug;7(8):1511–22.

82. Kumar M, Gouw M, Michael S, Sámano-Sánchez H, Pancsa R, Glavina J, et al. ELM—the eukaryotic linear motif resource in 2020. Nucleic Acids Res. 2020 Jan 8;48(D1):D296–306.

